# Cell type-specific CLIP reveals that NOVA regulates cytoskeleton interactions in motoneurons

**DOI:** 10.1101/237347

**Authors:** Yuan Yuan, Shirley Xie, Jennifer C. Darnell, Andrew J. Darnell, Yuhki Saito, Hemali Phatnani, Elisabeth Murphy, Chaolin Zhang, Tom Maniatis, Robert B. Darnell

## Abstract

**Background:** Alternative RNA processing plays an essential role in shaping cell identity and connectivity in the central nervous system (CNS). This is believed to involve differential regulation of RNA processing in various cell types. However, *in vivo* study of cell-type specific post-transcriptional regulation has been a challenge. Here, we developed a sensitive and stringent method combining genetics and CLIP (crosslinking and immunoprecipitation) to globally identify regulatory interactions between NOVA and RNA in the mouse spinal cord motoneurons (MNs).

**Results:** We developed a means of undertaking MN-specific CLIP to explore MN-specific protein-RNA interactions relative to studies of the whole spinal cord. This allowed us to pinpoint differential RNA regulation specific to MNs, revealing major role for NOVA in regulating cytoskeleton interactions in MNs. In particular, NOVA specifically promotes the palmitoylated isoform of a cytoskeleton protein Septin 8 in MNs, which enhances dendritic arborization.

**Conclusions:** Our study demonstrates that cell type-specific RNA regulation is important for fine-tuning motoneuron physiology, and highlights the value of defining RNA processing regulation at single cell type resolution.

## Background

A thorough understanding of the complexities of the mammalian CNS requires detailed knowledge of its cellular components at the molecular level. RNA regulation has a central role in establishing cell identity and function across the numerous cell types in the CNS [1–7]. Traditional whole tissue based methods are particularly limited in their power to delineate cell type-specific RNA regulation in the mammalian CNS due to its vast cellular diversity and architectural complexity. While separation or induction of specific cell types *in vitro* provides a practical way for cell type-specific analyses [1–3,8,9], alteration of cellular biology due to loss of physiological contexts presents a significant caveat to this approach.

Recent technological breakthroughs using RiboTag and BAC-TRAP mouse lines have allowed for translational profiling at single cell type resolution [10–13]. These studies revealed remarkable differences in the population of translating mRNAs across various CNS cell types, highlighting the degree of molecular heterogeneity among neuronal cells. These methods offer important ways to study translated mRNAs in specific cell types, but do not provide a way to define other types of cell type-specific RNA regulation. Here we develop a complimentary and more general means to study RNA processing and regulation in a cell type specific manner.

RNA processing is regulated by RNA-binding proteins. Two of the best-studied are NOVA1 and NOVA2, neuron-specific KH-type RNA binding proteins that bind to YCAY motifs and regulate alternative splicing and polyadenylation [14–16]. Using crosslinking and immunoprecipitation (CLIP), a method that allows stringent purification of protein-RNA complexes captured *in vivo*, we identified NOVA targets in mouse neocortex [16–19], and have estimated that NOVA participates in the regulation of ~7% of brain-specific alternative splicing events in mouse neocortex [20]. Interestingly, NOVA targets are specifically enriched for transcripts encoding proteins with synaptic functions - a group of transcripts that drives CNS cell type diversity [13,15].

Indeed, NOVA proteins are essential for the function of multiple neuronal cell types [21–24]. In particular, we previously uncovered a pivotal role for NOVA in maintaining spinal motoneuron (MN) survival and physiology [5,21,25]. Here, to more precisely define NOVA-regulated RNA processing in spinal MNs, we developed a new strategy combining BAC-transgenic mice and CLIP to identify MN-specific RNA regulation. NOVA targets in MNs were especially enriched for genes encoding microtubule-, tubulin- and cytoskeletal protein binding proteins. These results led us to uncover a NOVA-mediated RNA processing event differentially regulated in MNs involving a cytoskeleton protein, Septin 8, which controls dendritic complexity. Cell type-specific CLIP revealed a previously unidentified role of NOVA in motoneurons, highlighting the importance of cell-type specific analysis of RNA regulation.

## Results

### BAC transgenic mice express epitope tagged NOVA specifically in motoneurons

*Nova1* and *Nova2*, the two mammalian *Nova* paralogs, encode proteins harboring three nearly identical KH-type RNA binding domains. *Nova* genes are widely expressed among various neuronal cell types in the spinal cord, including the MNs. Our strategy to study MN-specific NOVA regulation is twofold – to express AcGFP (*Aequorea coerulescens* green fluorescent protein)-tagged NOVA specifically in MNs, followed by CLIP using antibodies against the AcGFP epitope tag.

To test if an N-terminal AcGFP tag alters NOVA RNA binding specificity, we compared global RNA binding profiles of NOVA2 and AcGFP-NOVA2. NIH/3T3 cells ectopically expressing NOVA2 or AcGFP-NOVA2 were subjected to HITS-CLIP using antibodies against NOVA and GFP, respectively (Figure S1A). We generated complex NOVA:RNA interactomes for both tagged and untagged NOVA2, as defined by 1,775,101 unique CLIP reads for NOVA2 and 9,778,818 for AcGFP-NOVA2. Both sets of CLIP data showed comparable genomic distributions (Figure S1B). We identified NOVA2 and AcGFP-NOVA2 CLIP peaks, defined as regions with significantly higher CLIP tag coverage than gene-specific background expected from uniform random distribution [26,27]. The canonical NOVA recognition motif, YCAY, was enriched around NOVA2 and AcGFP-tagged NOVA2 CLIP peaks to the same degree (Figure S1C-D; R^2^ = 0.94), with tetramers that may overlap YCAY by three or more nucleotides (YCAY, NYCA, CAYN) showing the highest enrichment at both NOVA2 and AcGFP-NOVA2 peaks (red dots, Figure S1D). These indicate that the N-terminal AcGFP tag does not perceptibly alter NOVA2 binding to its cognate RNA motifs.

Having confirmed the RNA binding fidelity of AcGFP tagged NOVA2, we generated transgenic mice expressing AcGFP-NOVA2 under the MN-specific choline acetyltransferase (*Chat*) promoter. The AcGFP-Nova2 cassette was inserted into a bacterial artificial chromosome (BAC) harboring the *Chat* promoter through homologous recombination (Figure 1A) [28]. The engineered BAC was injected into fertilized C57BL/6 oocytes and two Chat: GFP-Nova2 BAC transgenic lines, #6 and #17, were selected and maintained from five original founder lines.

**Figure1.**
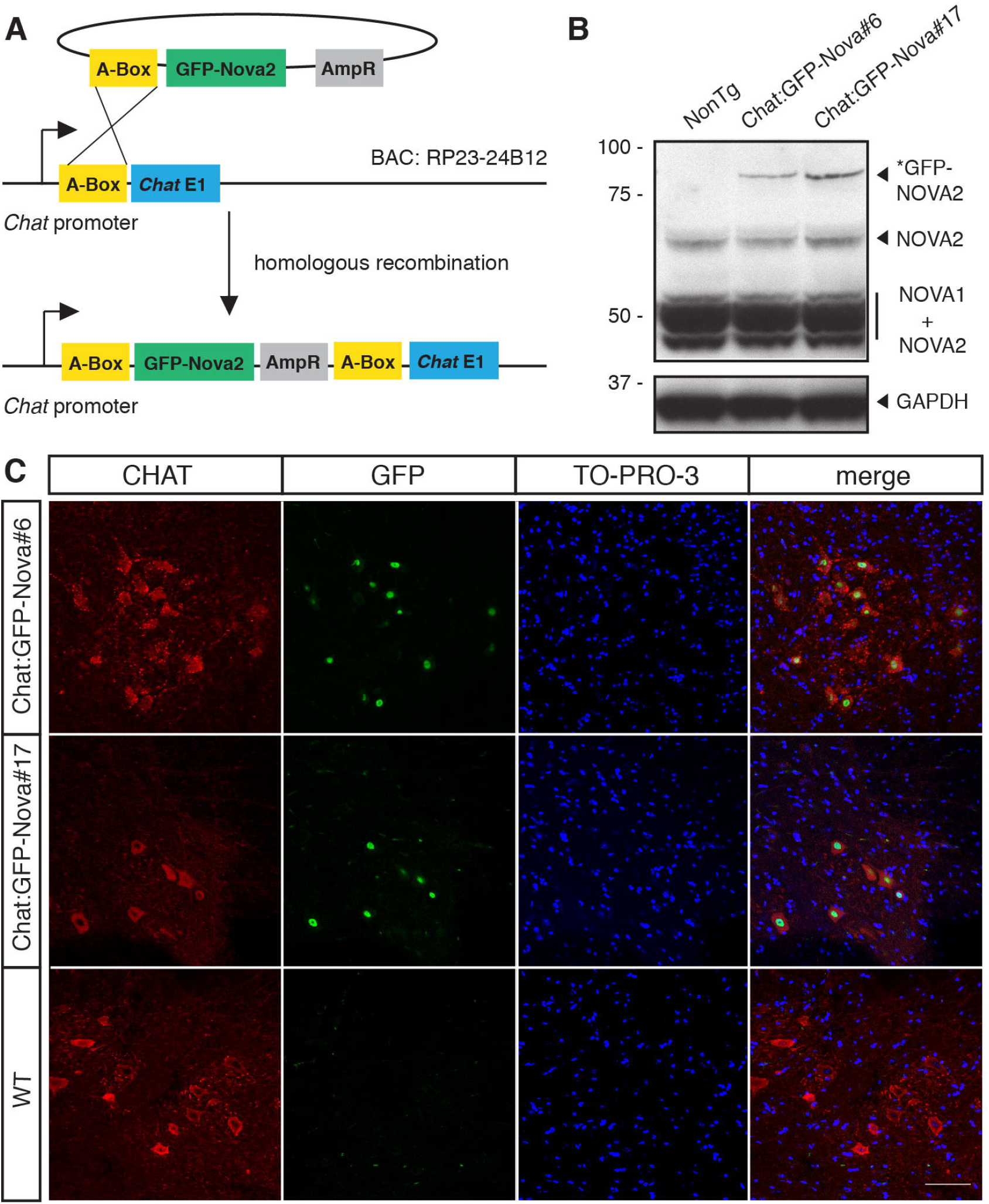
BAC-transgenic mouse lines express AcGFP-tagged NOVA2 in MNs. A. Schematic diagram illustrating the generation of recombinant BAC. A-box, a ~500 nt sequence homologous to the mouse *Chat* 5’-UTR region, mediated the insertion of a plasmid containing GFP-Nova2 coding sequence downstream of *Chat* promoter. B. Western blotting of spinal cord lysate from wild type (WT) mice and mice from Chat:GFP-Nova2 #6 and #17 lines using human anti-NOVA serum. The 93 kD GFP-NOVA2 was expressed only in transgenic spinal cords. GAPDH was blotted as a loading control. C. Immunofluorescence on spinal cord transverse sections using antibodies against CHAT and GFP, and counterstained with TO-PRO-3. Scale bar represents 50 μm.

Chat:GFP-Nova2 transgenic mice were born at expected Mendelian ratios and were phenotypically indistinguishable from wild type littermates throughout their lifespan (data not shown). Expression of the GFP-NOVA2 fusion protein was confirmed in the spinal cords of both Chat:GFP-Nova2 lines, but not the wild type controls (Figure 1B). To assess the MN-specific expression pattern of the transgene, we performed immunofluorescence staining on spinal cord transverse sections. GFP-NOVA2 expression was confined to CHAT-positive neurons in the transgenic lines (Figure 1C). Thus we have successfully established transgenic mouse lines with epitope tagged NOVA specifically expressed in MNs.

### HITS-CLIP generates a robust transcriptome-wide NOVA-RNA interaction map in motoneurons

To define NOVA binding sites in MNs, we undertook HITS-CLIP on Chat:GFP-Nova2 spinal cords using a mixture of two monoclonal antibodies against GFP to maximize avidity and specificity. GFP IP was performed using wild type or transgenic spinal cords with or without UV crosslinking, followed by ^32^P-labeling of bound RNA. The presence of labeled GFP-NOVA2-RNA complexes was dependent on both UV crosslinking and the expression from the Chat:GFP-Nova2 transgene, further demonstrating the specificity of GFP-NOVA2 CLIP under these conditions (Figure 2A, lanes 1-3). Partially digested RNA crosslinked to GFP-NOVA2 was subjected to subsequent library preparation steps (Figure 2A, lane 4, Methods). Importantly, the production of cDNA was dependent on reverse transcriptase (Figure 2B), confirming the absence of DNA contaminants in the CLIP RNA libraries. cDNA inserts between 30-80 nt (Figure 2B) were selected for next-generation sequencing (NGS) and further analysis.

**Figure 2.**
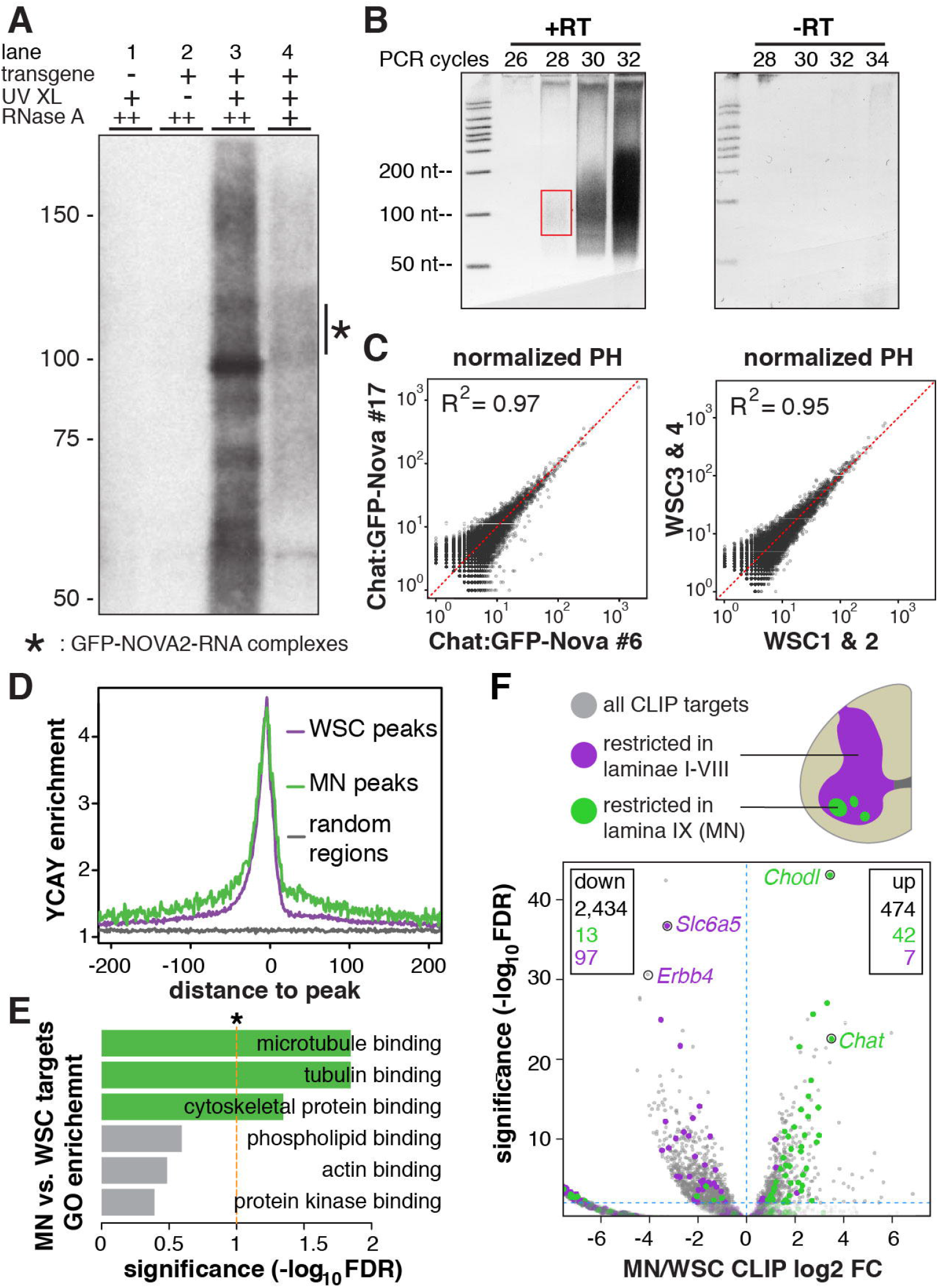
MN CLIP generated a *bona fide* and robust MN NOVA binding profile. A. Autoradiogram showing NuPAGE separation of radio-labeled GFP-NOVA2-RNA complexes. Both the transgene and UV crosslinking were required for the presence of radio-labeled GFP-NOVA2-RNA complex (lanes 1-3), which appeared as a smear from 100 to 125 kD with partial RNase digestion and collapsed to a band around 95 kD in high RNase concentration (lanes 3 and 4). B. Representative RT-PCR polyacrylamide gel images for CLIP library cloning. Left panel shows RT-PCR products at incremental PCR cycle numbers, and right panel shows control reactions without reverse transcriptase. The red box indicates the cDNA (with 5’ and 3’ linkers) size range purified for subsequent cloning and sequencing. C. Pairwise correlation of normalized CLIP peak height (PH) between #6 and #17 transgenic lines, and between two WSC groups. Normalized PH is the sum of raw CLIP reads in a given peak normalized to the read depth of each individual experiment. For comparison purpose, CLIP reads in WSC groups were randomly downsampled in the same number of peaks (27,628) to match the complexity of MN CLIP. D. Enrichment of YCAY around CLIP peaks. YCAY enrichment is calculated by normalizing the number of YCAY motifs starting at a given position relative to all WSC or MN CLIP peaks, to the expected YCAY frequency based on random base distribution. E. Molecular function GO enrichment of MN relative to WSC NOVA targets. FDR values were calculated using hypergeometic test, followed by Benjamini-Hochberg multiple test correction. The top six GO terms with the smallest FDR values are shown. Horizontal bars shows significance of GO enrichment, with significantly enriched GO terms in green, and nonenriched in grey. Dashed line marks the FDR value 0.1. F. Volcano plot of gene-wise CLIP reads enrichment in MN versus WSC. Illustration of spinal cord transverse section is shown on the top right corner, with Rexed laminae I-VIII in purple and lamina IX in green. Genes with a restricted expression pattern in Rexed lamina IX and laminae I-VIII, as curated by the Allen Spinal Cord Atlas, are shown as green and purple dots, respectively. All other NOVA CLIP targets are displayed as grey circles. Two MN and two interneuron marker genes are labeled. The numbers of NOVA CLIP targets with enriched or depleted NOVA binding in MN are indicated at the upper right and left corners of the chart, respectively, with font colors indicating the Allen Spinal Cord Atlas curated subgroups. The horizontal blue dashed line denotes FDR value 0.01.

Four sets of biological replicate HITS-CLIP experiments were performed on each Chat:GFP-Nova2 transgenic line, with each biological replicate consisting of pooled spinal cord samples from five to seven 3-month-old mice. We obtained a total of 2,023,726 unique CLIP reads from these 8 biological replicates. Since AcGFP-Nova2 expression is confined to MNs, we refer to this group of CLIP reads as MN NOVA CLIP reads. As a reference to all NOVA binding sites in the whole spinal cord, we additionally performed four standard NOVA CLIP experiments on endogenous NOVA in 3-month-old wild type mouse whole spinal cords (WSC), generating 17,353,049 unique WSC NOVA CLIP reads (Figure S2A, Additional file 1A). CLIP reads from MN CLIP showed higher intronic and less exonic distribution compared to those from WSC CLIP (p = 4.2 × 10^−7^, Figure S2B).

NOVA peaks in MN and WSC CLIP datasets were defined separately [27]. A total of 25,681 CLIP peaks (p ≤ 0.01) were identified for MN NOVA CLIP, which harbor CLIP reads originating from at least four of the eight biological replicates (biologic complexity (BC) ≥ 4 out of 8; see [16]; Additional file 1B). In parallel, we identified 218,794 WSC NOVA peaks represented in at least two out of four biological replicates (BC ≥ 2 out of 4, Additional file 1C). The CLIP data from our two transgenic lines was highly correlated (R^2^ = 0.97), as were WSC CLIP reads (Figure 2C), underscoring the ability of cell type-specific HITS-CLIP to reproducibly and quantitatively measure *in vivo* cellular protein-RNA interactions.

MN HITS-CLIP generated a genome-wide binding profile characteristic of endogenous NOVA proteins. The canonical NOVA binding motif YCAY was significantly and similarly enriched around MN and WSC CLIP peaks (Figure 2D). WSC NOVA CLIP peaks mapped to 12,445 mm9-annotated Entrez genes, including to the alternative spliced regions of 303 out of 335 known NOVA-regulated genes (Additional file 1E); MN CLIP peaks mapped to 4,450 Entrez genes, including to 174 known NOVA-regulated alternatively spliced regions (Additional file 1D) [20]. It is of note that when compared to all NOVA-regulated genes in the WSC, this subset of MN NOVA targets are especially enriched for genes encoding microtubule binding (GO:0008017), tubulin binding (GO:0015631) and cytoskeletal protein binding proteins (GO:0008092) (hypergeometric test, FDR < 0.05, Figure 2E and Additional file 1F), suggesting specialized functions for NOVA-RNA regulation in motoneurons.

We tested whether transcripts known to be enriched in MN, when compared with WSC, showed enrichment of NOVA binding in MN NOVA CLIP. The numbers of WSC or MN CLIP reads within respective peaks were summed for each gene, followed by analysis using the Bioconductor edgeR package [29] to identify genes with differential enrichment of CLIP reads in MN or WSC. 474 and 2,434 transcripts showed enriched and depleted NOVA binding in MN compared to WSC, respectively (FDR ≤ 0.01, Figure 2F and Additional file 1G & H). Consistent with the cell type specificity of GFP-NOVA2 CLIP, the known MN markers *Chat* and *Chod1* were among the top transcripts with the most significantly enriched GFP-NOVA2 CLIP reads coverage (FDR = 2.61 × 10^−23^, 8.39 × 10^−44^, respectively) [30] (Figure 2F). Conversely, *Slc6a5*, a glycine transporter gene known as an inhibitory neuron marker [31], as well as *Erbb4*, an interneuron-restricted receptor tyrosine kinase [32,33], were two transcripts with the most significantly depleted MN CLIP reads coverage (FDR = 1.84 × 10^−37^, 2.31 × 10^−31^, respectively, Figure 2F).

We further systematically examined transcripts with well-defined anatomic expression patterns in spinal cord. Spinal cord grey matter exhibits a pattern of lamination consisting of ten laminal layers, with large alpha-motoneuron pools located in lamina IX [34]. Based on the Allen Spinal Cord Atlas generated and curated in-situ hybridization data [35], 166 transcripts are exclusively expressed in lamina IX, of which 55 displayed differential NOVA binding (FDR ≤ 0.01) in MN. Meanwhile, 301 transcripts are restricted in one or more laminae other than lamina IX, of which 104 showed significant differential NOVA binding (FDR ≤ 0.01). Among this group of 159 (55 + 104) transcripts with differential NOVA binding in MN, the enrichment or depletion of NOVA CLIP signals on these transcripts in MN compared to WSC were highly concordant with transcript spatial expression patterns (green and purple dots in Figure 2F), in that 42 of the 55 (76%) lamina IX enriched transcripts had enriched NOVA binding in MN (p < 2.2 × 10^−16^, chi-squared test), and 97 out of the 104 (93%) lamina I-VIII enriched transcripts had lower NOVA binding in MN (p = 0.007, chi-squared test). Taken together, we conclude that HITS-CLIP in Chat:GFP-Nova2 lines captured MN-specific transcriptome-wide NOVA-RNA interactions.

### NOVA displays MN-specific binding patterns

We were particularly interested in NOVA binding sites along a given gene that were disproportionally strengthened or weakened in MNs compared to WSC (Figure S3). To identify such NOVA-RNA interactions, we bioinformatically pooled WSC and MN CLIP reads to define peak regions referred as Joint Peaks (JPs), and grouped these JPs by genes (Figure S3 and Methods). For each gene with two or more JPs, we performed pairwise Fisher’s exact test to identify JPs with disproportionally enriched MN or WSC CLIP reads (Figure S3). This subset of JPs denotes NOVA binding sites strengthened or weakened in MN compared to WSC.

Using this method, we identified 121,216 genic JPs after performing our peak-finding algorithm on pooled WSC and MN CLIP reads and filtering for biological reproducibility (Figure S3). Analysis of intronic and exonic JPs as two separate groups revealed 707 (1.07%) strengthened and 160 (0.24%) weakened intronic JPs in MNs (Figure 3A and Additional file 2A), as well as 1,058 (2.34%) strengthened and 407 (0.90%) weakened exonic JPs in MNs, (FDR ≤ 0.1, fold change ≥ 2) (Figure 3B and Figure S4). The AcGFP epitope tag does not significantly account for the observed NOVA binding differences between MN and WSC (Figure S4A-C, Figure 3A-B). Examples of disproportionally strengthened and weakened NOVA binding sites are shown in Figure 3C (intronic) and 3D (exonic).

**Figure 3.**
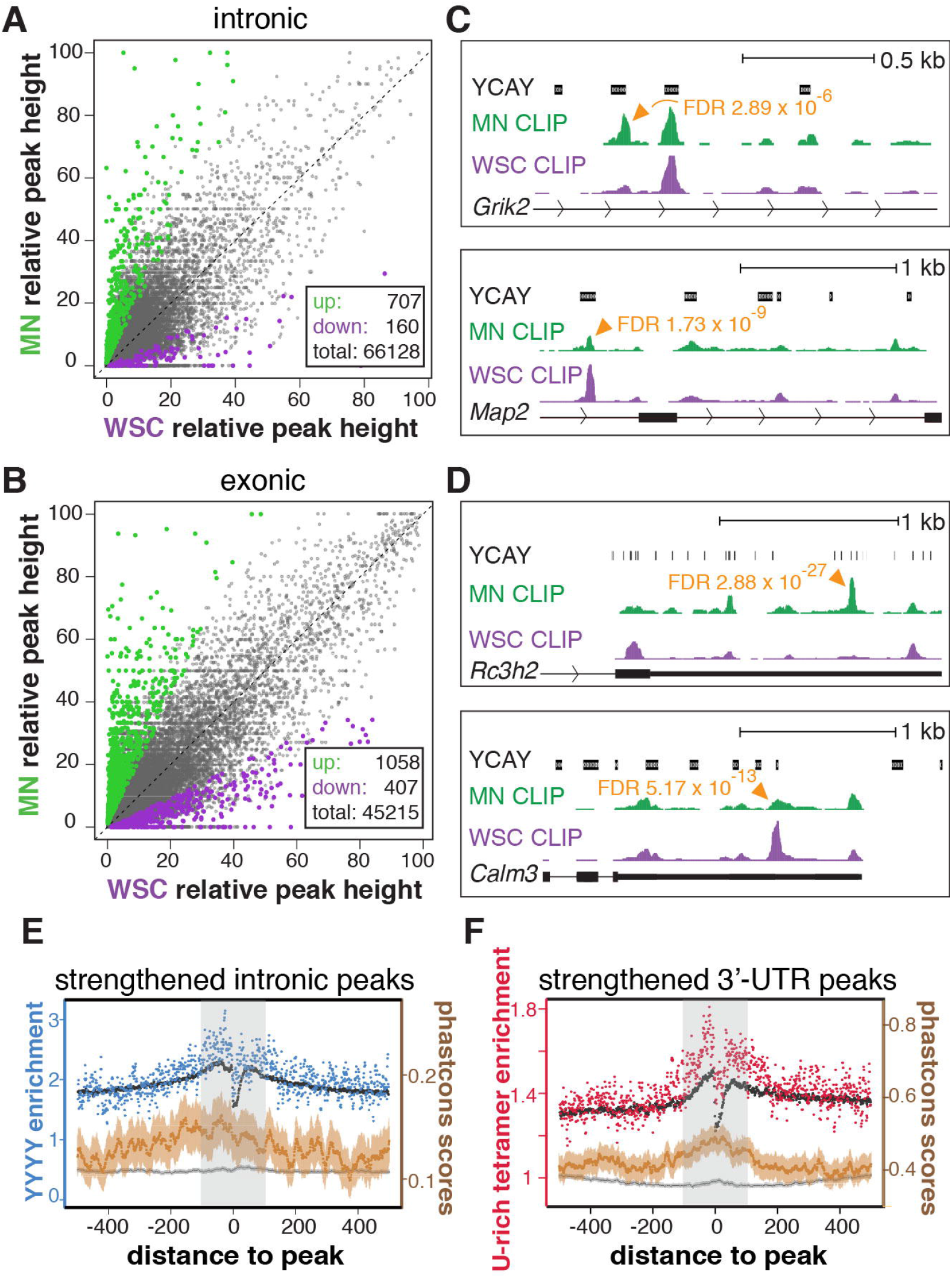
NOVA displays MN-specific binding patterns. A-B.Pairwise comparison of intronic and exonic relative NOVA peak heights in MN and WSC using the method illustrated in Figure S3. Relative NOVA peak heights in introns or exons are calculated as following: 100 * number of MN or WSC CLIP reads in a JP / number of MN or WSC CLIP reads in all intronic or exonic JPs in the corresponding gene. NOVA peaks disproportionally strengthened or weakened in MN (FDR ≤ 0.1, fold change ≥2) are shown in green and purple, respectively. The numbers of differential peaks and total peaks with sufficient coverage for the analysis are shown in the boxes. C-D. UCSC genome browser images illustrating disproportionally different intronic (C) an exonic (D) NOVA binding sites in MN. YCAY track demarcate clusters of NOVA binding motifs. The WSC and MN CLIP tracks are pooled HITS-CLIP results, normalized for the displayed regions so that the highest unchanged peaks in WSC and MN share the same height. Significantly different NOVA binding sites between MN and WSC are marked by arrowheads with FDR values indicated. UCSC gene annotation and transcript direction are shown at the bottom of each panel. E-F. Positional enrichment of YYYY and U-rich tetramers and phastcons scores around intronic (E) and 3’-UTR (F) NOVA peaks strengthened in MNs, respectively. YYYY and U-rich tetramer enrichment is calculated from motif frequencies at each base position relative to strengthened (blue or red) or all (black) MN CLIP peaks in introns or 3’-UTRs, normalized by their expected frequencies based on random base distribution. Phastcons scores are plotted with solid dots denoting the mean phastcons values at a given base position, and lighter lines denoting 95% confidence intervals. Dark grey and brown represent phastcons scores around all and strengthened intronic/3’UTR NOVA peaks, respectively. Motif enrichment scales are on the left, and phastcons score scales are on the right. Light grey boxes highlight regions 100 nt around NOVA peaks.

We examined potential RNA sequence signatures around MN-specific NOVA binding sites (FDR ≤ 0.1, fold change ≥ 2) by analyzing neighborhood tetramer distributions. We discovered a marked overrepresentation of polypyrimidine (YYYY) tetramers around intronic NOVA binding sites strengthened in MN, (Figure 3E), as well as a striking enrichment of U-rich (two or more uridines) tetramers around 3’-UTR NOVA binding sites strengthened in MN (Figure 3F). For both polypyrimidine and U-rich tetramers, differential enrichment is confined to regions 100 nt up- and downstream of strengthened MN peaks (Figure 3E-F). Interestingly, regions flanking changed MN peaks are evolutionarily more conserved compared to all intronic or 3’-UTR NOVA peaks (Figure 3E-F), suggesting the potential functional importance of differential NOVA binding sites in MN. Taken together, these data suggest that a large number of binding sites are differentially bound by NOVA independent of potential transcript level variations between WSC and MN, such that NOVA displays MN-specific binding patterns.

### MN-specific NOVA binding predicts MN-specific alternative splicing

Since NOVA plays an important role in regulating neuronal transcript splicing, we tested whether MN-specific NOVA binding correlates with MN-specific alternative splicing. We took advantage of a high quality motoneuron RNA-seq dataset, where spinal MNs in 3-month-old wild type mice were collected by laser capture microdissection (LCM) and subjected to RNA-seq [36]. For the WSC transcriptome, we performed RNA-seq on age, genotype and strain background matched whole spinal cords. Approximately 36 and 80 million mappable reads were obtained from either biological replicates of MN and WSC RNA-seq, respectively, and alternative splicing analysis was performed as described [26] (http://zhanglab.c2b2.columbia.edu/index.php/Quantas).

Around 30% (1,620 out of 5,418) of motoneuron expressed alternative exons showed biologically consistent differential splicing in MN compared to WSC (FDR ≤ 0.1, BC = 2 out of 2; Figure 4A and Additional file 3A). Strikingly, an even higher proportion of motoneuron expressed *Nova* targets are differentially spliced between MN and WSC (281 out of 626, 45%, FDR ≤ 0.1, BC2 out of 2, Figure 4A). The significant enrichment of *Nova* targets among exons differentially spliced in MNs (p < 2.2 x10^−16^, hypergeometric test) suggests that *Nova* contributes in an important way to MN-specific splicing patterns.

**Figure 4.**
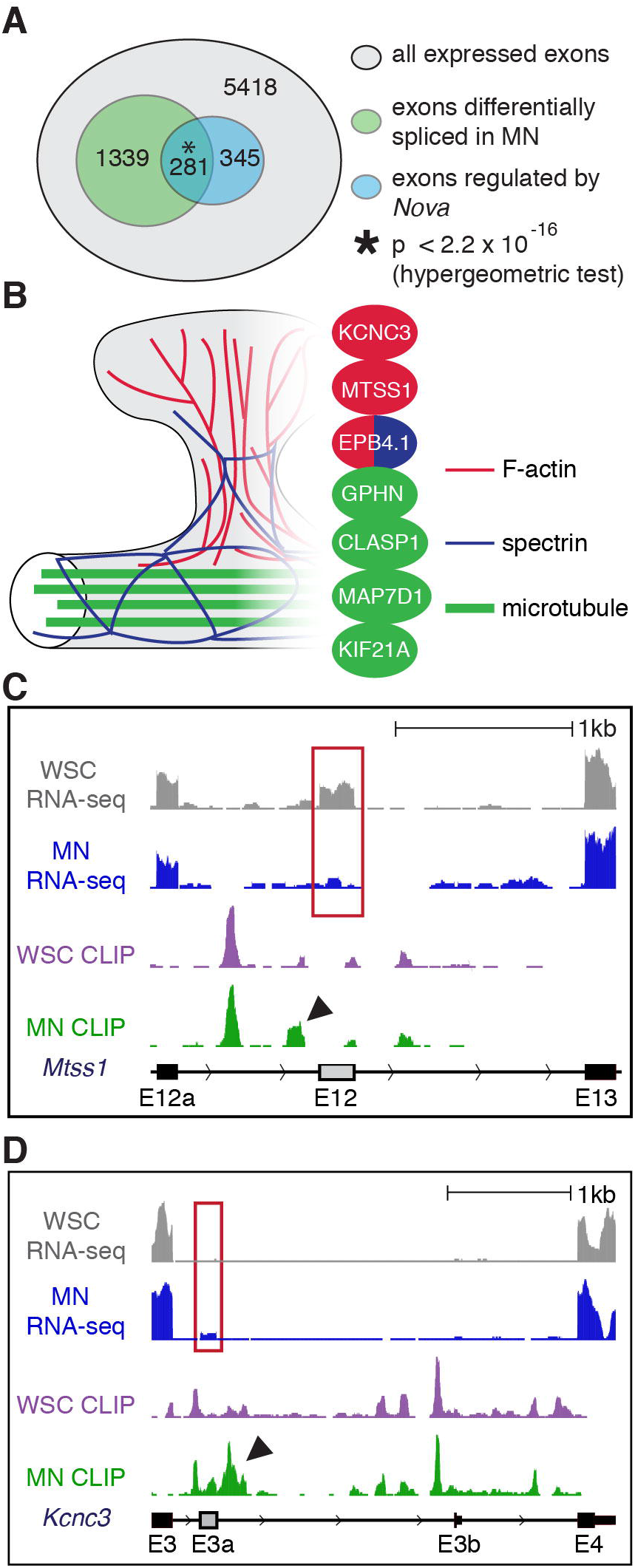
MN-specific NOVA binding correlates with MN-specific alternative splicing. A. Venn diagram showing overlap between cassette exons differentially spliced in MN and known *Nova* targets [20] among all expressed alternative exons (see Methods). P value is calculated by hypergeometric test. B. Illustration of cytoskeleton structures in part of a dendrite and a dendritic spine. Actin filaments are represented in red, microtubules in green, and spectrin in navy. Differentially regulated MN NOVA targets are represented in colors corresponding to their interacting cytoskeletal component(s). C-D.UCSC genome browser images illustrating correlation between differential NOVA binding and MN specific alternative splicing in the cases of *Mtss1* E12 (C) and *Kcnc3* E3a (D). YCAY track demarcate clusters of Nova binding motifs. WSC and MN RNA-seq tracks are RNA-seq results from 3-month-old whole spinal cord (this study) and laser dissected motoneurons [36], respectively, with biological replicates pooled. For alternative splicing visualization, these two tracks share the same maximum heights of flanking exons. Exons differentially spliced (FDR ≤ 0.1) in MN versus WSC are highlighted in red boxes. The WSC and MN CLIP tracks are pooled HITS-CLIP results, normalized for the given regions so that the highest unchanged peaks in WSC and MN share the same height. Significantly different NOVA binding sites between MN and WSC (FDR ≤ 0.1) are marked by arrowheads. UCSC gene annotation and transcript direction are shown at the bottom of each panel, with alternative exons marked in grey. For *Mtss1* E12, an increase of the NOVA peak (arrowhead) immediately upstream in MN correlates with increased E12b inclusion in MN. Similarly, a dramatic increase of NOVA binding immediately downstream of *Kcnc3* E3a (arrowhead) correlates with activated E3a splicing in MN.

We tested whether MN-specific NOVA binding correlated with MN-specific alternative splicing. Although NOVA binding in broader regions may influence splice site choice, high confidence predictions on whether NOVA promotes or represses alternative exons is achieved when NOVA binds within a window of 400 nt around the regulated exons [16] - upstream binding highly correlates with splicing repression, and downstream binding with splicing activation. Transcriptome wide, thirteen MN-strengthened or weakened NOVA binding sites (FDR ≤ 0.1) are located in these regulatory “hotspots” around alternative exons with RNA-seq coverage sufficient for analysis. Interestingly, the majority of these exons (9 out of 13, 69%) showed MN-specific splicing patterns (FDR ≤ 0.1). Based on our NOVA-RNA map, differential NOVA binding patterns in MNs compared to WSC correctly predicted increase or decrease of alternative exon inclusion in MNs compared to WSC for 8 out of the 9 alternative exons (89%) (Additional file 3B), consistent with NOVA mediating a direct action to regulate MN-specific alternative splicing.

Intriguingly, the seven genes hosting these nine MN-specific splicing events encode a functionally coherent set of proteins. Six of the seven gene products, i.e. MTSS1, MAP7D1, KIF21A, EPB4.1|3, CLASP1, GPHN, and KCNC3, are known to interact with cytoskeleton components (Figure 4B) [37–44]. For example, MTSS1 binds to actin monomers and induce membrane protrusion [37,45]. *Mtss1* harbors two alternatively spliced exons, E12 and E12a, at the 3’-end of its coding sequence, and inclusion of E12a, but not E12, is necessary to promotes neuritogenesis [46,47]. Interestingly, MN CLIP revealed a unique Nova binding site 109 nt upstream of *Mtss1* E12 in MN (3-fold increase in relative peak height; FDR = 0.046, Figure 4C), which would predict based on the NOVA-RNA map that NOVA binding would inhibit E12 in MN. Indeed, we observed a remarkably lower E12 inclusion rate in MN compared to WSC (dI (MN-WSC) = −0.69, FDR = 9.65 × 10^−7^; Figure 4C).

Another example is *Kcnc3*, which encodes a pan-neuronal voltage-gated potassium channel Kv3.3 [48,49]. MN CLIP and RNA-seq revealed a new alternative exon E3a which showed a significantly higher inclusion rate in MNs compared to WSC (dI (MN-WSC) = 0.05, FDR = 1.38 × 10^−14^). The increased utilization of E3a in MNs positively correlated with a dramatically enhanced NOVA binding site in MN 121 nt downstream of E3a (4-fold increase, FDR = 4.02 × 10^−7^) (Figure 4D). Interestingly, inclusion of E3a would lead to a Kv3.3 isoform with an extended C-terminal proline-rich domain, which has been shown to modulate channel inactivation through triggering actin nucleation at the plasma membrane [43]. Taken together, these observations suggest that unique NOVA binding patterns around alternative exons in MN contributes to MN-specific biology, particularly in shaping the cytoskeleton and regulating cytoskeleton interactions in unique ways within spinal cord MNs.

### NOVA differentially regulates *Sept8* alternative last exon usage in MNs

Regulation of alternative last exon (ALE) usage involves intricate interplay between splicing and cleavage/polyadenylation, two processes both regulated by NOVA [16]. We therefore investigated the potential role that NOVA played in regulating ALE usage in MNs. 65 ALEs in 34 genes were differentially included in MNs compare to WSC (FDR ≤ 0.1, |dI| ≥ 0.2, Figure 5A and Additional file 4A), with the top differentially utilized ALEs residing in two functionally related genes, *Sept8* and *Cdc42* (Figure 5A) [50–52]. We also examined the published *Nova2* WT and KO mouse brain RNA-seq dataset [24], and identified additional NOVA regulation on 17 ALEs in 9 genes (FDR ≤ 0.1, |dI| ≥ 0.2, Figure 5B and Additional file 4B). Interestingly, among all genes with ALEs, *Sept8* was the only NOVA target differentially regulated in MN compared to WSC.

**Figure 5.**
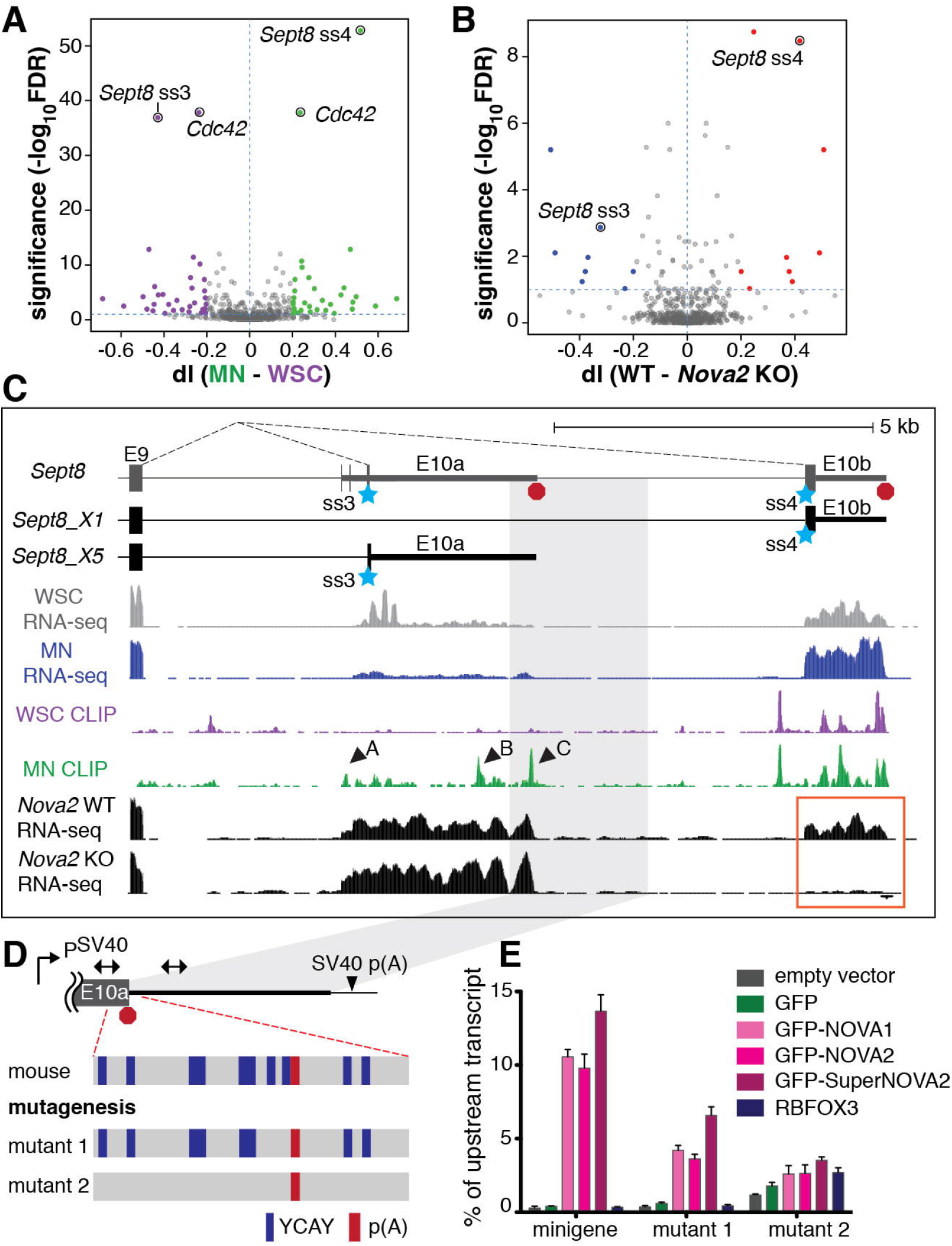
NOVA promotes *Sept8* exon 10b usage in MNs. A-B.Volcano plot of differential ALE usage in MN versus WSC (A) and *Nova2* WT versus KO mouse brains (B). ALEs with higher inclusion rates in MNs and *Nova2* WT (dI ≥ 0.2, FDR ≤ 0.1) are labeled in green and red, respectively. ALEs with lower inclusion rates in MNs and *Nova2* WT (dI ≤ 0.2, FDR ≤ 0.1) are labeled in purple and blue, respectively. The blue horizontal dashed lines denote FDR value 0.1. C. UCSC genome browser images illustrating the correlation between differential NOVA binding pattern and *Sept8* ALE usage in MN. Partial gene and transcript structures of *Sept8* are shown on top. Alternative last exons 10a and 10b are utilized in the X5 and X1 isoforms, respectively. Blue stars mark the two predominant alternative 3’ splice sites used in the adult spinal cord. Red octagons mark polyadenylation sites. For WSC and MN RNA-seq and NOVA CLIP tracks, see Figure 4 C-D legend for reference. Arrowheads mark significantly strengthened NOVA binding sites in MN. E18.5 WT and *Nova2* knockout mouse brain RNA-seq are displayed in black. Inclusion of exon 10b is dependent on NOVA, as highlighted by the orange box. Light grey box marks the genomic region included in the Sept8 minigene in D. D. Illustration of the Sept8 minigene construct. The 3’-end of *Sept8* exon 10a and its adjacent intronic sequence was inserted downstream of the SV40 promoter, and upstream of the SV40 poly(A) signal. Double-arrowed segments represent qPCR amplicons used for measuring transcription readthrough. The region surrounding the poly(A) signal is enlarged, with YCAY motifs marked in navy, and the poly(A) signal in red. Mutant 1 minigene lacks the two YCAY motifs proximal to the poly(A) signal. Mutant 2 minigene is devoid of YCAY in the 150 nt region surrounding the poly(A) signal. E. Sept8 minigene assay. COS-1 cells expressing minigene variants and indicated proteins were harvested for RNA isolation and qRT-PCR analysis. Two amplicons illustrated in Figure 5D were used to measure RNA levels up- and downstream of exon 10a poly(A) sites, respectively. Y-axis represents percentage of downstream relative to upstream transcript level. Error bars represent standard error of the mean (SEM) based on three independent replicates.

*Sept8* encodes a family member of the septin proteins, which are multi-functional components of the cytoskeleton [53,54]. In neurons, septins regulate dendritic and axon morphology through modulating actin and microtubule dynamics [55–57]. Although much is known about other septins, SEPT8 is a more recently described family member and less well characterized. Mouse *Sept8* harbors two alternative terminal exons, exon 10a and 10b (Figure 5C). Four mutually exclusive splice acceptors in *Sept8* exons 10a and 10b can directly join downstream of exon 9, with splice acceptors 3 in exon 10a and 4 in exon 10b utilized in >85% of *Sept8* transcripts in adult mouse spinal cords (Figure 5C). Comparison of ALE usage altered between *Nova2* WT and KO mouse brains [24] showed markedly lower inclusion rate for exon 10b in *Nova2* KO (dI = 0.42, FDR = 3.32 × 10^−9^; Figure 5C), suggesting that NOVA promotes splice acceptor 4 usage, exon 10a exclusion and exon 10b inclusion (Figure 5C). In MNs, RNA-Seq data indicate that the majority of *Sept8* transcripts use splice acceptor 4 (65% in MN vs. 22% in WSC), while splice acceptor 3 is preferentially bypassed (20% in MN vs. 72% in WSC, Figure 5C). This observed differential ALE usage between MN and WSC coincides with higher NOVA binding in MNs at three binding sites located in exon 10a (site A: fold change = 10, FDR = 0.0050; site B: fold change = 3.4, FDR = 0.0046; site C: fold change = 4.4, FDR = 3.84 × 10^−5^, Figure 5C). NOVA binding in site C is in a highly conserved YCAY-dense region encompassing the putative polyadenylation site at the 3’-end of exon 10a (Figure 5D, Figure S5A).

NOVA has been shown to prevent cleavage/polyadenylation through binding in close proximity to polyadenylation sites [16]. We tested whether NOVA association with the 3’-end of *Sept8* exon 10a blocked cleavage/polyadenylation, thus allowing for RNA polymerase II (PolII) readthrough and inclusion of the downstream exon 10b. A minigene was constructed using the genomic region of the 3’-end of exon 10a and the 5’ part of intron 10 (Figure 5D). We co-transfected this minigene reporter with vectors expressing NOVA proteins or controls into COS-1 cells (Figure S5B), which lack endogenous NOVA, followed by quantitation of RNA levels up- and downstream of the E10a polyadenylation site. In the absence of NOVA proteins, transcription terminated efficiently at the E10a polyadenylation site, as measured by the less than 0.5% transcription readthrough rate (Figure 5E). Exogenous NOVA proteins increased the downstream readthrough dramatically (>10%, >25 fold, Figure 5E), while another RNA-binding protein (RBFOX3) showed no effect on cleavage/polyadenylation at E10a (Figure 5E), indicating that NOVA proteins efficiently blocked cleavage/polyadenylation and promoted downstream transcription.

To test whether direct NOVA association with the YCAY sites is necessary for the observed regulation, we generated two mutant minigenes by disrupting YCAY sites while preserving the GC content (Figure 5D). Mutant1, with the two YCAY sites proximal to the E10a poly(A) site mutated, showed moderately dampened responses (6-11 fold) to NOVA overexpression (Figure 5E). In contrast, little NOVA regulation (< 2 fold, Figure 5E) was observed for mutant 2, where all YCAY sites within 150 nt of the poly(A) site were disrupted. Taken together, these results suggest that NOVA promotes *Sept8* exon 10b inclusion by binding close to the polyadenylation site in exon 10a and boosting read-through transcription and utilization of exon 10b, and that strengthened NOVA binding around exon 10a poly(A) site in MNs lead to higher exon 10b usage observed in MNs.

### The *Sept8* exon specifically promoted by NOVA in MNs encodes a palmitoylated filopodia inducing motif (FIM) that enhances dendritic arborization

Alternative usage of exon 10a and 10b confers the C-terminal variation between SEPT8 protein isoforms X5 and X1, respectively. SEPT8 undergoes palmitoylation in vivo [58], which is a reversible post-translational process of attaching a 16-carbon saturated fatty acid to cysteine residues [59]. Although the definitive palmitoylation site(s) in SEPT8 is unknown, the only predicted sites are two exon 10b encoded cysteine residues (C469, C470) in the NOVA-promoted SEPT8-X1 isoform_[60]. To assess whether C469 and C470 mediate SEPT8-X1 palmitoylation, we expressed SEPT8 isoforms in COS-1 cells and employed the acyl-resin-assisted-capture (acyl-RAC) assay for palmitoylation detection [61,62]. This assay relies on the indirect capture of palmitoylated proteins facilitated by palmitoyl-ester specific cleavage (Figure S6A). Whereas acyl-RAC failed to detect any palmitoylated SEPT8-X5, 5-10% of SEPT8-X1 was shown to be palmitoylated as evidenced by the presence of SEPT8-X1 in the resin captured fraction (Figure 6A, lane 2). This capture was dependent on palmitoyl ester specific cleavage (Figure 6A, lanes 3 & 4). Mutations in C469 and C470 (SEPT8-X1-mut, C469S/C470S) prevented the detection of SEPT8-X1 palmitoylation (Figure 6A), indicating that the two cysteines encoded by the NOVA-promoted exon 10b are the sites for palmitoyl addition. The data here collectively suggests that NOVA-regulated alternative RNA-processing in MNs mediates isoform-specific SEPT8 palmitoylation.

**Figure 6.**
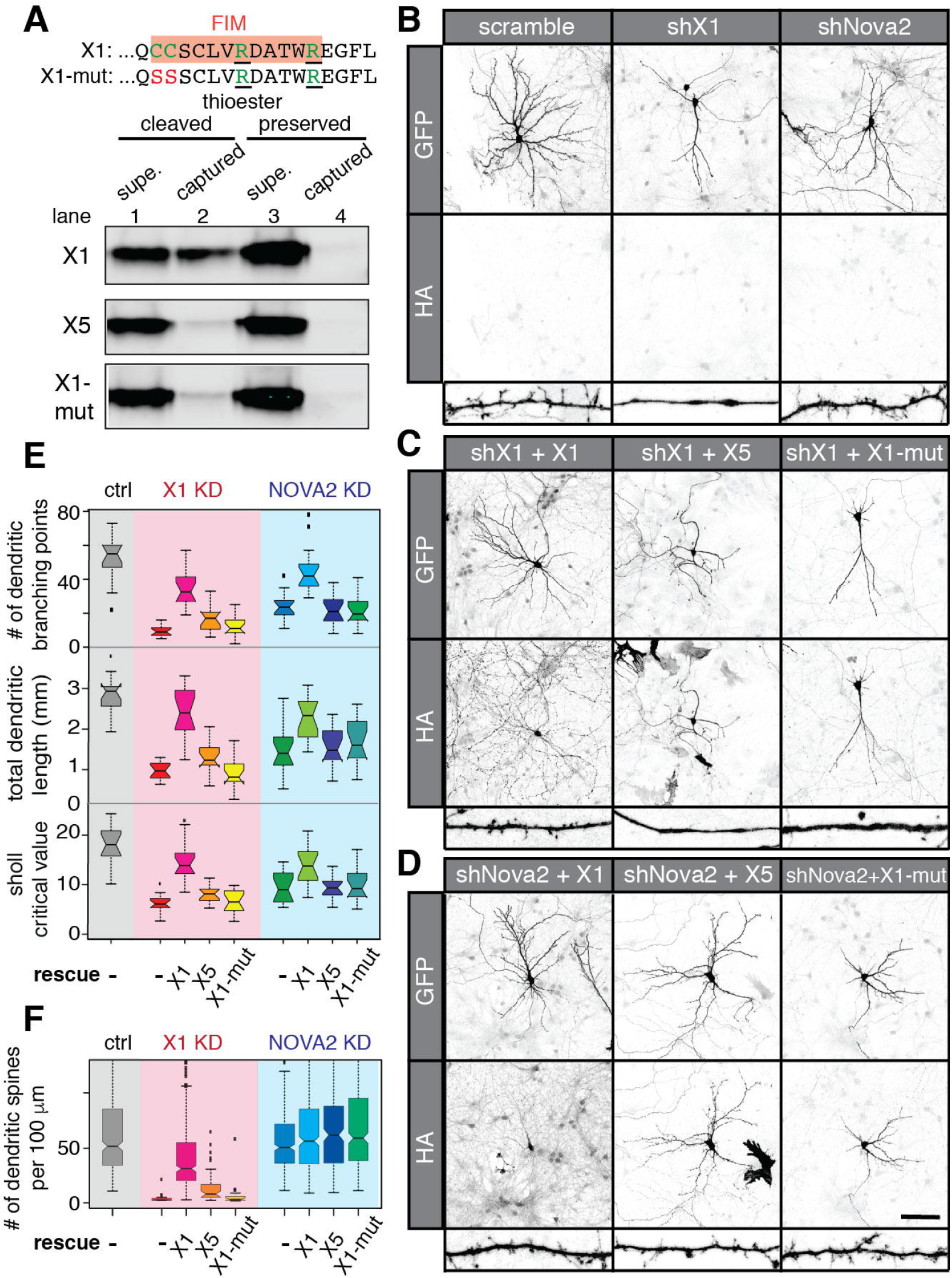
NOVA-dependent SEPT8 isoform promotes dendritic arborization and spine morphogenesis. A. Detection of palmitoylated SEPT8-X1 by acyl-RAC (as illustrated in Figure S6A). Top: C-terminal amino acid sequence of SEPT8-X1. Orange box highlights the FIM motif, with green letters marking palmitoylated cysteines and nearby basic amino acid residues. C469 and C470 were mutated to serine residues in SEPT8-X1-mut. Bottom: COS-1 cells expressing HA-tagged SEPT8 variants were used for the acyl-RAC assay. Immunoblotting of supernatant and thiopropyl-sepharose captured fractions using an antibody against HA is shown. 10% of supernatants were loaded compared to the captured fractions. B-D. Representative maximum projected confocal images of hippocampal neurons expressing shRNAs and HA tagged SEPT8 or controls. GFP, which is co-expressed from the shRNA constructs, is used as a neurite tracer. Anti-HA labels exogenously expressed SEPT8 variants. Representatively images of dendrites are shown at the bottom of each panel. Scale bar represents 100 μm. E-F. Boxplots evaluating dendritic arbor complexity (E) and dendritic spine density (F) in neurons with SEPT8-X1 or NOVA2 KD and SEPT8 rescue. Number of dendritic branching points, total dendritic lengths, sholl analysis critical values are plotted in E, and number of dendritic spines per 100 μm is plotted in F. Measurements in neurons expressing the control, shX1, and shNova2 shRNAs are highlighted by the grey, pink, and blue boxes, respectively. Quantification was based on 15-17 neurons per group.

Interestingly, we discovered that *Sept8* exon 10b encodes a potential filopodia inducing motif (FIM), characterized by two adjacent palmitoylated cysteines (C469, C470) and nearby basic residues (R475, R480) (Figure 6A, orange box) [63]. It has been shown that the palmitoylated FIM motif was sufficient to promote dendritic branching and spine formation in neurons [63]. To assay the functional effects of SEPT8-X1 on dendritic morphology, we performed isoform-specific knockdown (KD) using a construct co-expressing GFP and an shRNA targeting *Sept8* exon 10b (shX1) in primary mouse hippocampal neurons which express SEPT8-X1 (Figure S6B). KD efficiency (79%) and isoform specificity of shX1 was demonstrated in COS-1 cells expressing exogenous SEPT8-X1 and X5 (Figure S6C). Compared to neurons expressing the control scramble shRNA, neurons transfected with shX1 had significantly less complex dendritic arbors, as indicated by an 82% reduction in the number of dendritic branching points (Figure 6B & 6E, p = 2.9 × 10^−9^). Meanwhile, total dendritic length showed a 65% reduction (p = 9.2 × 10^−11^), and sholl analysis revealed a similar 66% reduction in critical value in neurons with SEPT8-X1 KD (Figure 6E, p = 4.4 × 10^−10^). Even more strikingly, SEPT8-X1 KD led to an almost complete absence of dendritic spines – dendrites became smooth with a few focal swellings at distal ends (Figure 6B & F, p =7.0 × 10^−82^). These abnormalities were partially rescued by co-expression of the shRNA-resistant SEPT8-X1, but not SEPT8-X5 or the palmitoylation deficient SEPT8-X1-mut (Figure 6C, E-F). Therefore, we conclude that SEPT8-X1 promotes dendritic branching and spine formation through its palmitoylated FIM motif.

Since NOVA promotes the SEPT8-X1 isoform, we predicted that neurons lacking NOVA would exhibit a similar reduction in dendrite arbor and spine density. We knocked down NOVA2 using an shRNA (shNova2) which efficiently depleted 76% of the endogenous NOVA2 in N2A cells (Figure S6D). Similar to SEPT8-X1 knockdown, neurons depleted of NOVA2 displayed greatly decreased dendritic arbors compared to control shRNA transfected neurons – as evidenced by a 55% reduction in the number of dendritic branching points, a 47% reduction in total dendritic length, and a 48% reduction in the sholl analysis critical value (Figure 6B & E, p = 3.6 × 10^−7^, 2.5 × 10^−7^, 1.8 × 10^−7^, respectively). On the other hand, to our surprise, NOVA2 knockdown did not significantly affect dendritic spine density (Figure 6B & F). Co-expression of SEPT8-X1, but not the two non-palmitoylated variants, partially restored dendritic arbor complexity in neurons with NOVA2 KD (Figure 6D-E), suggesting that NOVA2 enhances dendritic arborization through promoting the FIM-containing SEPT8 isoform.

## Discussion

Understanding brain function involves understanding its parts, as demonstrated by the discovery of differences in ribosome-associated transcripts evident by looking at specific neuronal cell types in the basal ganglia (D1 vs D2 neurons) [12]. Here we develop a general method combining BAC-transgenic engineering and CLIP that is conceptually applicable to the study of any protein-RNA interactions within specific cell types. We apply this strategy to analyze differential NOVA binding in mouse spinal motoneurons, compare that to interactions visible at the gross level of whole spinal cord analysis, and uncover NOVA-regulated biology specific to motoneurons. MN-specific CLIP revealed a major role for NOVA in regulating cytoskeleton interactions in MN, a function obscured in previous whole tissue-based analyses. This led us to discover a consequent defect in dendritic morphology in neurons lacking NOVA, which may help explain in part the severe MN defect seen in NOVA1/2 double KO motoneurons [21].

### MN specific CLIP unmasks previously undetected aspects of NOVA function

Motoneurons are among the largest neurons in the central nervous system. Their distinct morphology, characterized by a long axon and intricate dendritic arbor, renders traditional cell purifications based on enzymatic tissue digestion and cell purification methods particularly unsatisfactory for understand the molecular biology of the whole neuron, particularly given abundant evidence for RNA localization within the dendritic arbor. The cell type-specific CLIP strategy developed here allows robust and quantitative identification of NOVA binding sites in motoneurons *in vivo*, in both the cell bodies and processes. This generated a transcriptome-wide NOVA-RNA interaction atlas in motoneurons with NOVA binding sites in over 4,000 genes. NOVA binding in motoneurons reflects MN transcriptome signatures, which further demonstrated the cell type specificity of our assay.

The cell-type specific CLIP strategy developed here allowed delineation of a subset of MN NOVA targets from the whole spinal cord. These MN NOVA targets are enriched in genes encoding synaptic functions to a similar extent compared to WSC NOVA targets, yet they are especially enriched in genes encoding cytoskeleton interacting proteins. Furthermore, through combining cell type-specific CLIP with MN transcriptome profiling, we discovered RNA processing events differentially regulated by NOVA in motoneurons. Around 2% of NOVA binding sites were strengthened or weakened in MNs, which is consistent with our findings that ~2% (8 alternatively splicing regions out of 303) of NOVA regulated alternative splicing showed MN-specific splicing patterns correctly predicted by MN-specific NOVA binding. Interestingly, the vast majority of these events also reside in transcripts encoding cytoskeleton-interacting proteins. These findings reveal a previously undiscovered cell-type specific role for NOVA in regulating MN cell biology.

### NOVA plays an important role in regulating MN cytoskeleton interactions

The top NOVA-regulated ALE was in *Sept8*, which encodes a member of the multifunctional septin family that is capable of regulating neurite outgrowth and branching through interactions with cytoskeleton components [55–57]. NOVA binding at the polyadenylation site in *Sept8* exon 10a correlated with exon 10b usage, and this was only evident in analysis of MN CLIP, not WSC CLIP alone. Consistent with prior observations of position-dependent effects of NOVA on APA [16], we demonstrated that NOVA directly inhibits cleavage/polyadenylation by binding close to the exon 10a poly(A) site, presumably promoting exon 10b transcription and splicing. Interestingly, the NOVA-dependent exon 10b encodes an FIM motif capable of promoting dendritic branching and spine formation. Indeed, we discovered that SEPT8-X1, the NOVA-dependent SEPT8 isoform harboring the FIM, specifically promotes dendritic arborization and spine formation. Moreover, we found that neurons lacking NOVA2 displayed decreased dendritic arbors, which are partially rescued by SEPT8-X1. Taken together, these data indicate that NOVA promotes dendritic arborization through *Sept8* regulation.

While SEPT8-X1 promotes dendritic spine formation, we were not able to detect changes in dendritic spine density in NOVA2 KD neurons. This may be related to insufficient effects from NOVA2 KD, or from contributing indirect factors, such as NOVA promotion of protein isoforms with antagonistic effects on spine formation. For example, based on in-silico palmitoylation site prediction [60], we identified a total of eight NOVA targets where a predicted FIM motif is regulated by NOVA. Of these eight genes, NOVA promotes the FIM-harboring isoform in four (*Sept8, Ccp110, Sdccag3*, and *Ccdc84*), while inhibiting the FIM encoding exon in the other four (*Sgce, Clip1, Kcnma1*, and *Ube2e2*). It is possible that changes of various NOVA targets upon NOVA KD mitigate the overall effect on dendritic spine density.

We have previously shown that NOVA proteins play an essential role in motoneuron physiology [21]. Motoneurons in mice lacking both *Nova* family members were paralyzed and failed to cluster acetyl-choline receptors at the neuromuscular junctions [21]. Successful rescue of the NMJ defect was evident after Nova-knockout motoneurons were engineered to constitutively express the *Nova*-regulated Z+ alternatively spliced isoform of agrin, but surprisingly remained paralyzed. Phrenic nerve stimulation revealed that the axonal synaptic machinery for conducting action potentials and synaptic vesicle release from the axon was functional, leading to the conclusion that dysregulation of additional motoneuron NOVA RNA targets contribute to a proximal physiologic defect in *Nova*-knockout motoneurons. Here MN-specific CLIP reveals a major role of NOVA in regulating the motoneuron cytoskeleton, including promoting dendritic complexity. Dendrites are the main information receiving sites of neurons. Spinal motoneurons, as the gateway controllers of the CNS motor outputs, have elaborate dendritic structures to meet the highly complex demand of precisely coordinating muscle contractions spatially and temporally [64,65]. Early and progressive dendritic degeneration has been reported in lower MNs in motoneuron disease mouse models as well as amyotrophic lateral sclerosis (ALS) patients [66,67], suggesting the importance of dendritic integrity to motoneuron function. Our new findings may provide additional avenues for understanding the role that NOVA and RNA regulation plays in motoneuron function.

### Differential NOVA binding sites in MN suggests combinatorial control of multiple RNA-binding proteins

When we compared the NOVA-RNA interactions in MN and WSC, we uncovered over 2,000 sites transcriptome-wide that are differentially bound by NOVA in MN compared to WSC. In this way cell-specific CLIP offers the possibility of revealing cell-type specific regulatory phenomenon that are otherwise obscured from analysis of whole tissues. What underlies this unique NOVA binding profile in motoneurons? It has been shown that combinatorial control is integral in RNA processing regulation. Cellular RNA-binding protein networks play a pivotal role in determining target selection and binding dynamics of a given RNA-binding protein [68]. For NOVA, interactions with PTBP2, the neuronal polypyrimidine tract binding protein, as well as RBFOX proteins have previously been demonstrated [20,69]. In particular, we have shown that PTBP2 interacts with and antagonizes NOVA in the regulation of glycine receptor alpha 2 subunit (*Glra2*) splicing [69]. Interestingly, compared to all NOVA binding sites, sequences around NOVA bindings sites that are strengthened in MN showed enrichment of pyrimidine-rich motifs and higher evolutionary conservation (Figure 3E-F). This sequence signature around MN-specific NOVA binding sites along with lower PTBP2 levels in MN [36] suggests the hypothesis that lower abundance of PTBP2 in MN allows for unmasking of certain NOVA binding sites that are otherwise occupied by PTBP2 in other neurons. This kind of intricate cell type specific interaction networks may help determine *Nova* target selection beyond the specificity determined by sequence and structural constraints.

### Cell type-specific CLIP

Cell type-specific CLIP is in general applicable to any cell type and a great variety of RNA-binding proteins. The current study is neuron-specific due to nature of *Nova* [25,70], but a further array of cell types within the CNS or other tissues can be studied *in vivo* through epitope-tagging. Indeed, Schaefer and colleagues used a similar strategy to identify AGO-bound miRNAs in D2 neurons of the mouse striatum [71], and *Camk2a*-neurons in the forebrain [72]. And as CLIP is applied to identify functional protein-RNA interactions defining the actions of an increasing number of RNA-binding proteins including splicing factors [19], RNA-editing factors [73], and epigenetic regulators [74], expanding aspects of post-transcriptional and RNA-related regulation can be investigated in a cell-type specific manner. We recently described a method, PAPERCLIP, in which PABPC1 CLIP was used to identify brain-specific transcripts and alternative 3’ UTR processing within those cells [75]. Combining a cell-specific tagging strategy with PAPERCLIP would allow profiling of cell-specific polyadenylated transcripts in the brain or within any tissue, an approach that would compliment the delineation of ribosome associated transcripts demarcated by BAC-TRAP.

We examined whether GFP-NOVA2 expression in our BAC-transgenic motoneurons could have affected our observations relative to unperturbed neurons. The BAC-transgenic system has intrinsic expression biases, and might, for example, have skewed the motoneuron NOVA-PTBP ratios, impacting the observed NOVA binding increases around pyrimidine-rich regions. However, the high concordance between NOVA binding changes in our transgenic motoneurons and differential splicing in MN without exogenous NOVA strongly argues for physiological relevance of our transgenic model. A more elegant approach of studying cell type specific RNA-protein interactions would be to generate conditionally epitope-tagged knock-in lines followed by the introduction of a cell type specific Cre recombinase expression. Epitope-tagged protein would be expressed in the desired cell type from its endogenous promoter, thus preserving protein stoichiometry.

Cell type-specific CLIP may be relevant for the study of many neurological diseases, such as ALS and ataxias, where defined cell types are pathologically affected during the entire course or the initial stages of disease progression. Interplay between the affected cells and their cellular context plays an important role in disease progression [76–79]. Our strategy allows for *in vivo* dissection of both cell autonomous and non-autonomous effects resulting from disease-causing mutations or insults, which had not be possible before. Great insights on cell type contributions to disease pathogenesis will be gained upon application of this strategy to a variety of animal models.

## Conclusions

Here we demonstrate the feasibility and physiological relevance of delineating neuronal cell type-specific RNA regulation. Through cell type-specific epitope-tagging of the RNA binding protein NOVA2, we generated a motoneuron-specific NOVA-RNA interaction map using CLIP. Cell type-specific CLIP revealed a major role of NOVA in regulating cytoskeleton interactions in motoneurons, including promoting the palmitoylated isoform of a cytoskeleton protein, Septin 8, which enhances dendritic arbor complexity. MN-specific NOVA binding predicts MN-specific alternative RNA processing, further supporting the idea that cell type-specific RNA regulation contributes to cell identity. Our study highlights a non-incremental gain of knowledge moving from whole tissue-to single cell type-based RNA regulation analysis. As our strategy for cell type-specific CLIP is highly adaptable, we envision wide application of this method in a variety CNS cell types as well as disease models.

## Methods

### Antibodies and dilutions

The following antibodies were used in CLIP experiments: anti-GFP mouse monoclonal antibodies 19F7 and 19C8 (Memorial Sloan Kettering Monoclonal Antibody Facility), anti-NOVA serum from a paraneoplastic opsoclonus-myoclonus ataxia (POMA) patient. The following antibodies and dilutions were used for immunoblotting: anti-NOVA patient serum (1:2000), mouse anti-GAPDH monoclonal antibody 6C5 (Abcam, 1:20,000), rabbit anti-RBFOX3 (1:500) [80], rabbit anti-HA monoclonal antibody C29F4 (Cell Signaling Technology, 1:100). The following antibodies and dilutions were used for immunofluorescence: rat anti-GFP monoclonal antibody (nacalai tesque, 1:1000), goat anti-GFP polyclonal antibody (Rockland, 1:500), goat anti-CHAT polyclonal antibody (EMD Millipore, 1:500), rabbit anti-HA monoclonal antibody C29F4 (Cell Signaling Technology, 1:1000), rabbit anti-SEPT8 monoclonal antibody (pan SEPT8, EPR16099, 1:200), mouse anti-SEPT8_X1 monoclonal antibody D-11 (Santa Cruz, 1:50), chicken anti-MAP2 antibody (Thermo Fisher Scientific, 1:1000).

### Cell culture and transfection

NIH3T3 and COS-1 cells were cultured in Dulbecco’s modified Eagle’s medium (DMEM) supplemented with 10% heat-inactivated fetal bovine serum, 100 U/mL penicillin and 100 μg/mL streptomycin. NIH3T3 and COS-1 cells were transfected with plasmid constructs using lipofectamine 2000 (Invitrogen), according to the manufacturer’s instructions.

Mouse hippocampal neurons were isolated from 18-day-old CD1 mouse embryos and cultured in 24-well plates according to established protocols. For shRNA knockdown and rescue experiments, 0.4 μg of shRNA vector and 0.6 μg of protein expressing vector or control were transfected at DIV 10 using 3 μL of NeuroMag following manufacturer’s protocol. Transfected neurons were fixed at DIV 12 for immunofluorescence staining and con-focal imaging (see below).

### Construction of BAC-transgenic mouse lines expressing GFP-NOVA2 in motoneurons

In brief, AcGFP-fused *Nova2* coding sequencing was cloned into pLD53.SC2 plasmid. Subsequently, a DNA fragment homologous to *Chat* 5’-UTR region was PCR amplified using primers GCCAGGCATCTGAGAGGC and CCTAGCGATTCTTAATCCAGAGTAGCAGAGCTG and inserted into the plasmid through blunt end ligation at AgeI site. The sequence of the resulting plasmid pLD53.SC2-AcGFP-Nova2 was confirmed by Sanger sequencing and deposited to the Addgene database. Recombinant BAC was generated using RP23-246B12 and pSC2-GFP-Nova2 as described previously [81], and microinjected into pronuclear oocytes of C57BL/6 mice from Charles River. By PCR genotyping, nine mice from 68 offspring were confirmed to carry the transgene. Two founders, #6 and #17, were bred with C57BL/6 mice from Charles River to establish stable transgenic lines.

### Immunofluorescence staining

PFA fixed spinal cords were transversely sectioned at 14 μm thickness, and kept at −80 °C. Before immunofluorescence staining, sections were rehydrated in PBS for 10 minutes, followed by an one-hour block in blocking buffer containing 100 mM Tris-HCl, pH 7.5, 150 mM NaCl, 0.2% Triton X-100, 5% horse serum. Primary antibodies diluted in blocking buffer were subsequently added to the sections followed by overnight incubation at RT. Secondary antibody hybridization and TO-PRO-3 staining was performed using Alexa Fluor conjugated antibodies diluted in 100 mM Tris-HCl, pH 7.5, 150 mM NaCl, 0.2% Triton X-100 with 1 μM TO-PRO-3. Confocal images were taken using inverted LSM 510 laser scanning confocal microscope (Zeiss).

For IF staining on hippocampal neurons, cells were fixed in 4% PFA at room temperature for 10 mintues, permeablized with 0.1% Triton-X 100 in PBS at room temperature for 15 mintues, and blocked in PBS containing 0.1% Tween-20 and 10% donkey serum for one hour at room temperature. The primary antibodies, diluted in PBS containing 1% BSA and 0.1% Tween-20, were added to the cells and incubated overnight at room temperature. Alexa Fluor conjugated secondary antibodies (Jackson Immunoresearch) were used at 1:500 dilution to detect either the tag epitope or proteins of interest. Confocal images were taken using inverted LSM 880 NLO laser scanning confocal microscope (Zeiss).

### HITS-CLIP experiments and analysis

HITS-CLIP experiments were performed as previously described [16,26] with PCR primers specified in Additional file 5. For motoneuron specific CLIP, 400 μL of Dynabeads protein G (Invitrogen) precoated with 50 ug of each anti-GFP monoclonal antibody (mAb) was used to immunoprecipitate GFP-NOVA2 in UV crosslinked spinal cord lysate pooled from five to seven 3-month-old BAC-transgenic mice. For WSC CLIP, 400 μL of Dynabeads protein G precoated with 80 μL of human anti-NOVA serum was used to immunoprecipitate lysate from one 3-month-old wild type mouse spinal cord. cDNA libraries were prepared as previously described and sequenced on Illumina Genome Analyzer IIx, HiSeq 1000 or HiSeq 2000.

Bioinformatic processing and mapping of CLIP NGS reads was performed similarly as previously described with slight modifications [18,80,82,83]. Specifically, we removed 3’ linker sequence from CLIP reads before mapping the remaining sequences to the mm9 build of mouse genome with novoalign (http://www.novocraft.com) without iterative trimming. Following mapping, CLIP reads immediately upstream of genomic GTGTC and several highly similar pentamers were removed. We have recently discovered that under our reverse transcription (RT) conditions, 3’ end of the RT primer could prime reverse transcription by hybridizing to GUGUC or similar pentamers in the CLIP’ed RNA, leading to their preferential amplification (Park C, personal communication). Bioinformatically removing such reads circumvents this technical caveat.

NOVA binding peaks were defined as previously described using gene regions compiled from refseq, UCSC known genes and ESTs plus their downstream 10 kb as transcription units [26,27,80]. Gene regions with Entrez IDs were referred to as known genes. Defined peaks were then filtered for biological complexity, requiring CLIP reads from at least half of the biological replicates. Joint Peaks (JPs), in particular, were defined by running the peak finding algorithm using CLIP reads pooled from both comparison groups (e.g., MN and WSC), and requiring CLIP reads from at least half of the biological replicates in either group (i.e., WSC BC 2 out of 4 or MN BC 4 out of 8).

Gene-wise CLIP reads enrichment between WSC and MN was evaluated using the Bioconductor edgeR package with tagwise dispersion model [29]. P-values of differential NOVA binding on specific sites were calculated using fisher’s exact test on the following 2 × 2 matrix:

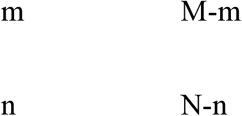

m and n stand for the number of MN and WSC CLIP reads in a given JP, respectively, while M and N stand for the total number of intronic/exonic MN and WSC CLIP reads within all intronic/exonic JPs in the corresponding transcript, respectively. Coverage is defined as the smallest number among the 2×2 matrix. Benjamini-Hochberg multiple-testing correction was performed using all intronic or exonic JPs with a minimum coverage of 10.

### RNA-seq library preparation and data analysis

We performed WSC RNA-seq on two biological replicates of 3-month-old SOD1^WT^ transgenic mice in SJL/B6 mixed background. For each replicate, poly(A) selected RNA from spinal cord was used to prepare RNA-seq library using Illumina TruSeq RNA sample prep kit. 100 nt paired-end reads were generated using Illumina Genome Analyzer II. In order to compare WSC and MN RNA-seq, we trimmed our WSC reads to 74 nt to match the read length of the MN RNA-seq dataset by Bandyopadhyay et al (GSE38820). Both WSC and MN datasets were mapped to the mouse (mm9) genome and exon junctions using OLego with default parameters (15 nt seed with 1 nt overlapping, ≤ 4 mismatches per read) [84]. Only reads unambiguously mapped to the genome or exon junctions were retained for downstream analysis. For MN RNA-seq, 36,484,398 and 38,293,208 NGS reads were mapped for the two biological replicates, respectively, while 74,444,847 and 81,175,598 NGS reads were mapped for WSC RNA-seq replicates. Transcript level and alternative splicing analyses were performed as previously described. Bioconductor edgeR package was used to evaluate statistical significance of transcript level differences between WSC and MN [29]. Fisher’s exact test was used to calculate p values of differential alternative splicing, and the FDR was estimated using the Benjamini-Hochberg method. Alternative exon splicing was analyzed as described [26]. Alternative exons with splice junction read coverage over 10 [26] were considered “expressed”, and were included in Benjamini-Hochberg multiple test correction. Differential alternative splicing events were identified by requiring FDR ≤ 0.1 in addition to biological consistency (BC2 out of 2).

### Gene ontology analysis

Gene ontology (GO) analysis was performed using GOrilla by running unranked lists of target and background genes [85]. For background genes, we used all genes with rpkm ≥1 in WSC or MN.

### Motif enrichment analysis

For motif enrichment around differential MN intronic NOVA peaks, tetramer occurences 100 nt around the center of each peak were counted, and compared to those 100 nt around all intronic or exonic NOVA peaks using hypergeometric test.

### Sept8 minigene assay

To construct the *Sept8* minigene, C57BL/6 genomic region amplified with primers TACGACTCACTATAGGGCGAATTCGGATCCGCATGAATTCTGACCCCTGTGA and CAATAAACAAGTTCTGCTTTAATAAGATCTCCGTAACCTGGCTACCAGTGA was cloned into BamHI and BglII digested pSG5 vector using HIFI DNA assembly kit (New England Biolabs). Mutant minigenes were constructed by assembling the PCR amplified vector backbone (forward: TTGGGCAGTAGCTTCGCTG, reverse: TAACATAACAGAGAAGCAAGCTGGCT) with synthetic gBlocks (IDT DNA) carrying the desired mutations. 1 μg of minigene was co-transfected into COS-1 cells with 1μg of control vector or NOVA/RBFOX3 expression vector. 48 hours post-transfection, cells were harvested for immunoblotting or RT-qPCR following standard protocols. Primers CTGACCCCGCATATGTTCCTGTGT and CAAGCTGGCTATCCTGGGCCTCTT were used to amplify an exon 10a region, while primers GTCTGCGATGGTTTTGCAGAGGTG and GGCCACAGGAAATGGAGATGTGAG were used for an intron 10 amplicon. Three independent experiments were performed for statistical comparison.

### Acyl-RAC assay

250,000 COS-1 cells were seeded per well in 6-well plates, and transfected with 2 ug of constructs expressing FLAG-HA tagged SEPT8 variants 18-24 hours later. Acyl-RAC assay was performed using the CAPUREome S-Palmitoylated Protein Kit (Badrilla) according to the manufacturer’s protocol. Treated protein lysates were subsequently immunoblotted with rabbit anti-HA antibody.

### Image analysis

Maximum projected confocal images were produced using ImageJ using only linear adjustments. Dendrites were semi-automatically traced using the Simple Neurite Tracer plugin. Critical value and number of dendritic branching points were deduced by using the “sholl analysis” and “analyze skeleton” plugins. Fifteen to seventeen neurons were analyzed per group.

## Declarations

### Ethics approval and consent to participate

All animal experiments were performed in the Association for Assessment and Accreditation of Laboratory Animal Care (AAALAC) – accredited Animal Resource Center at the Rockefeller University under protocol numbers 07111, 14678, and 17013.

### Consent for publication

Not applicable

### Availability of data and material

Raw .fastq files of RNA-seq and HITS-CLIP data have been deposited to the NCBI Gene Expression Omnibus (GEO) under accession number GSE71294.

### Competing interests

The authors declare that no competing interests exist.

### Funding

This work was supported by the National Institutes of Health grant 5RC2NS069473-02. R.B.D. is a Howard Hughes Medical Institute Investigator.

### Authors’ contributions

RBD conceived the project. YY, SX, JCD, AJD, YS, HP, and EM conducted experiments. YY and CZ analyzed data. YY and RBD wrote the manuscript with feedback from all co-authors. RBD and TM provided resources and supervision. All authors read and approved the final manuscript.

## Acknowledgements

The authors would like to thank Rada Norinsky, Roxana Cubias for oocyte microinjection, Nathaniel Heintz for pLD53.SC2 plasmid, Michael Moore, Christopher Park, Mariko Kobayashi for helpful discussion, and Scott Dwell for high-throughput sequencing.

---

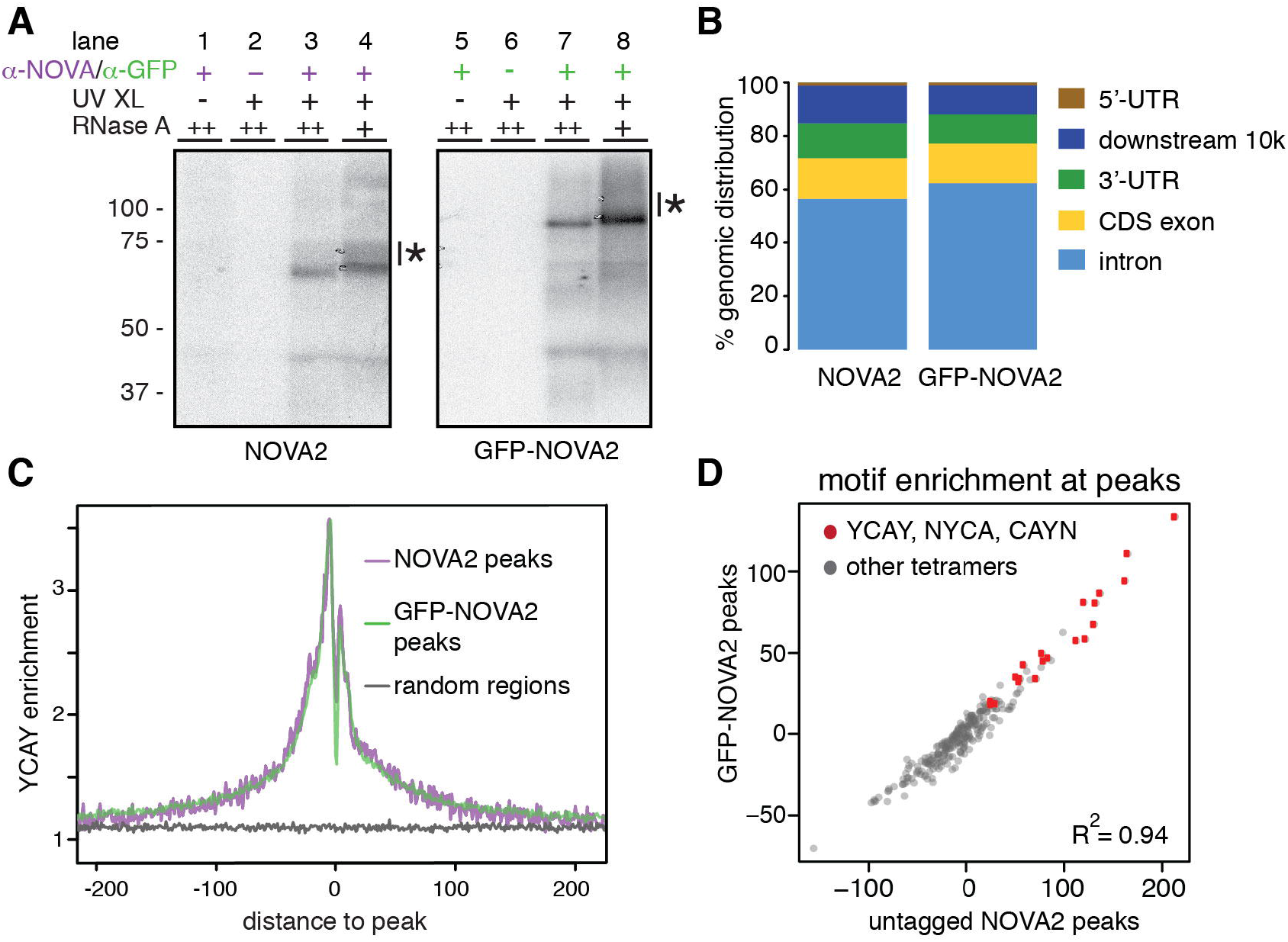

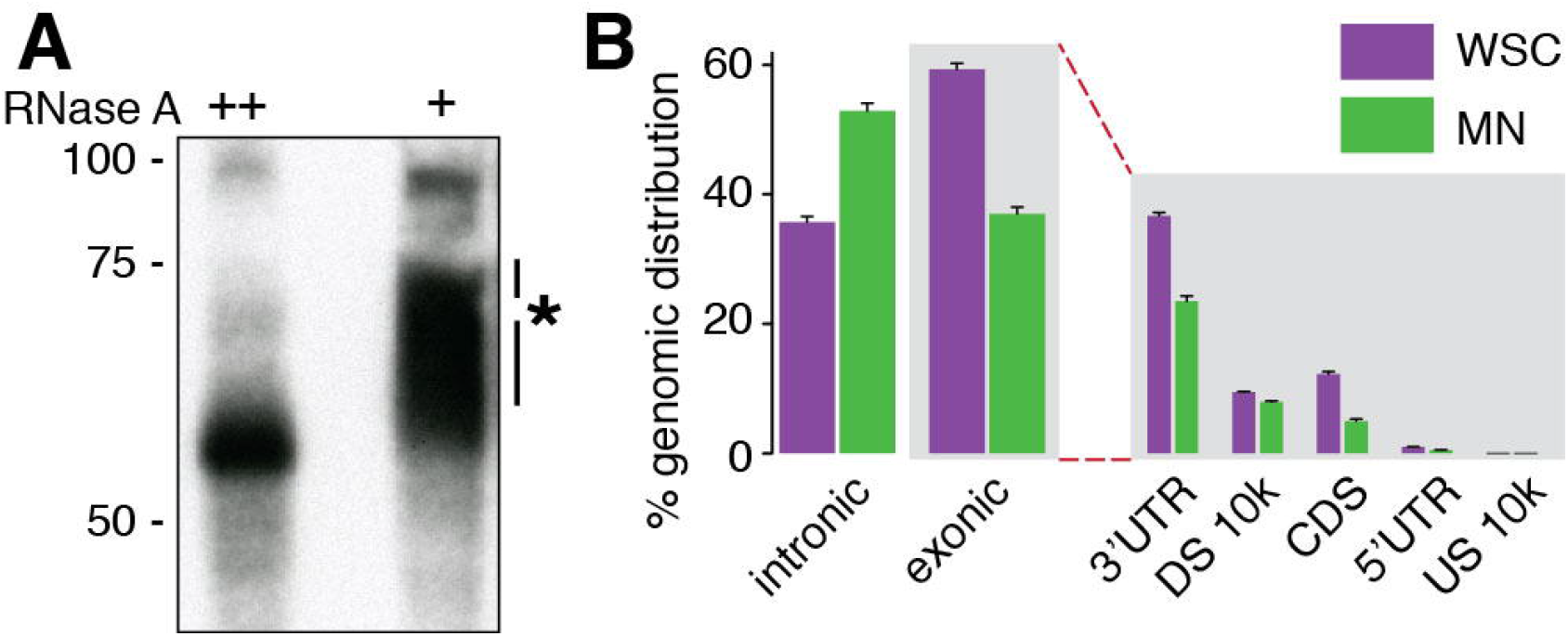

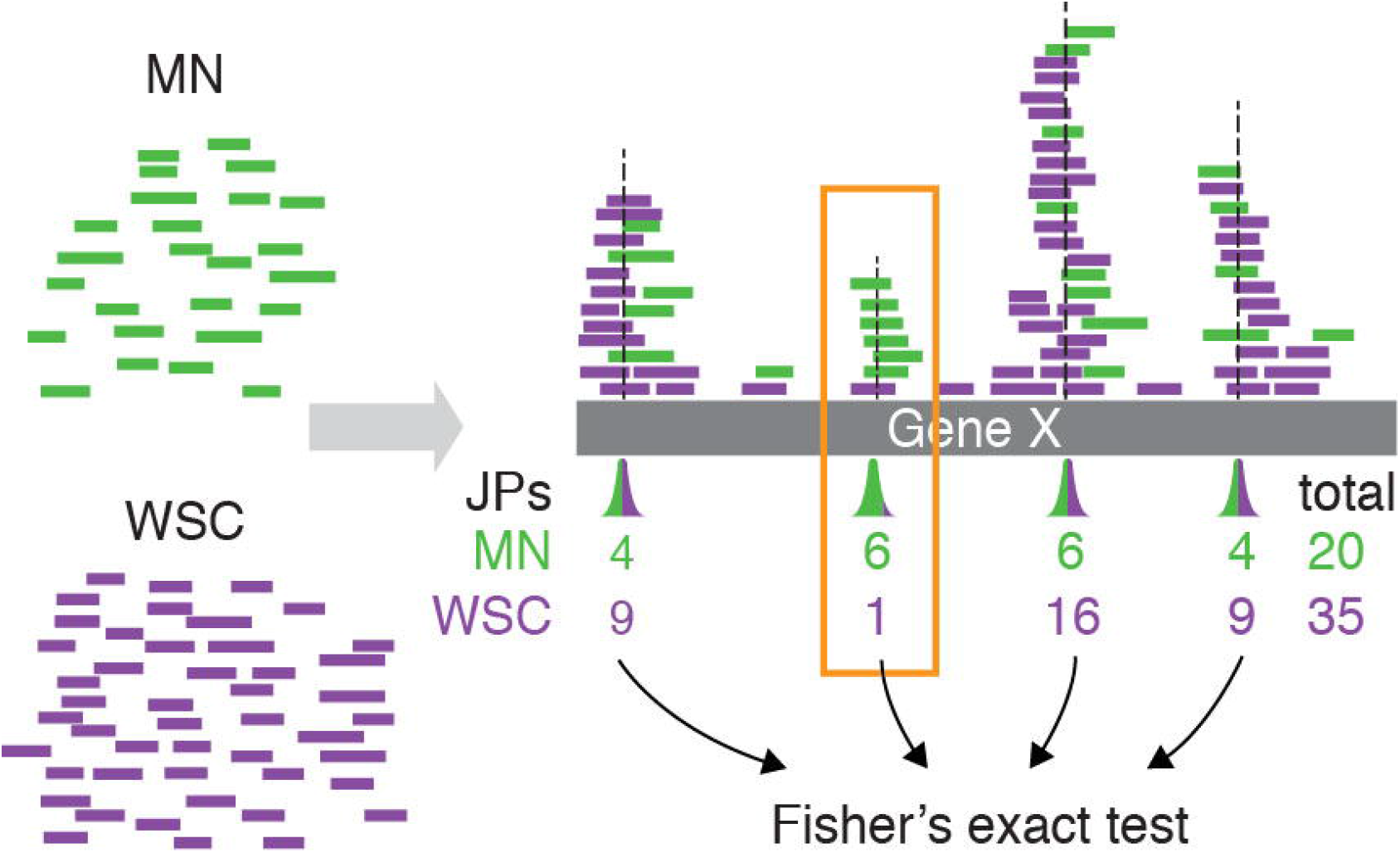

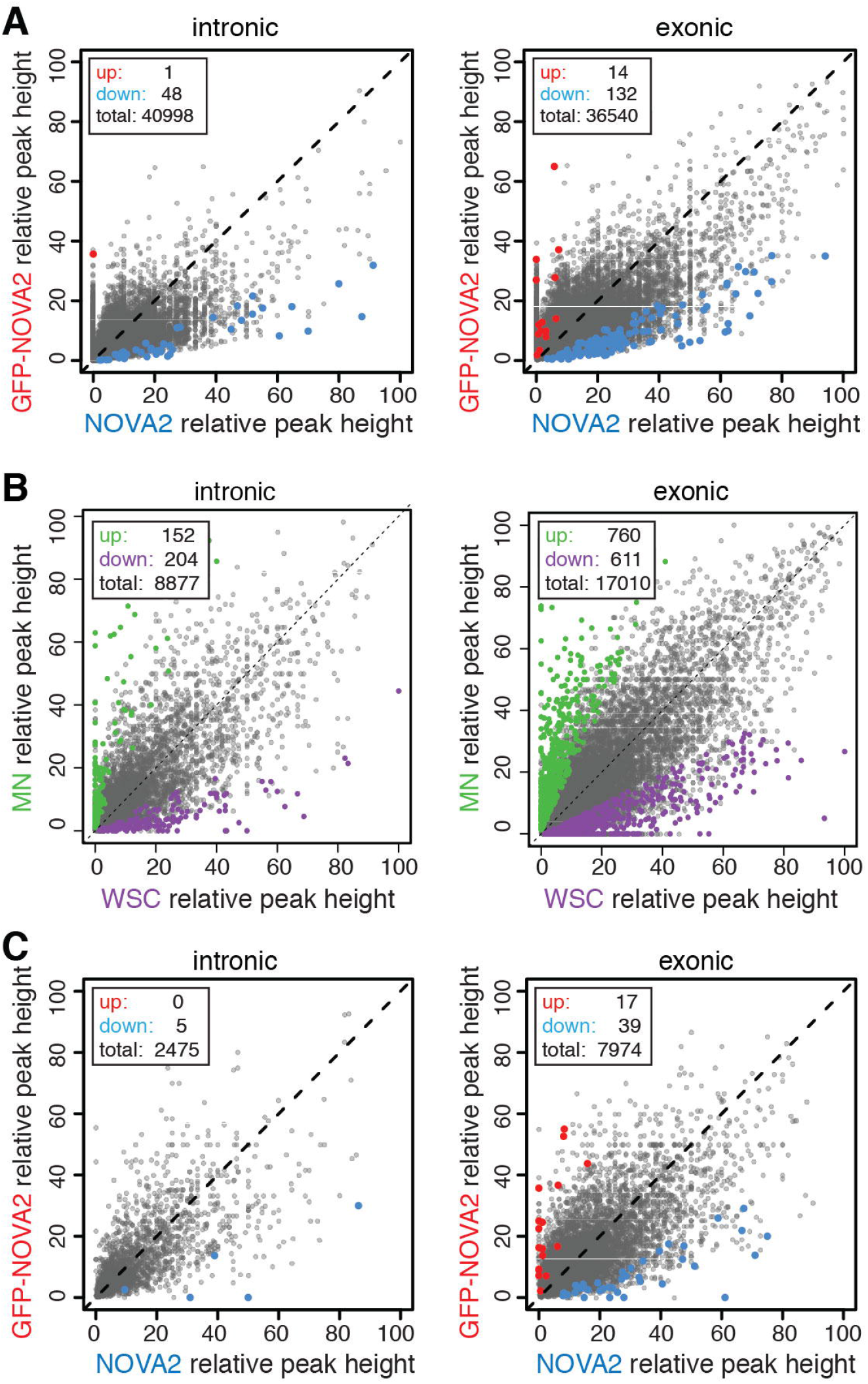

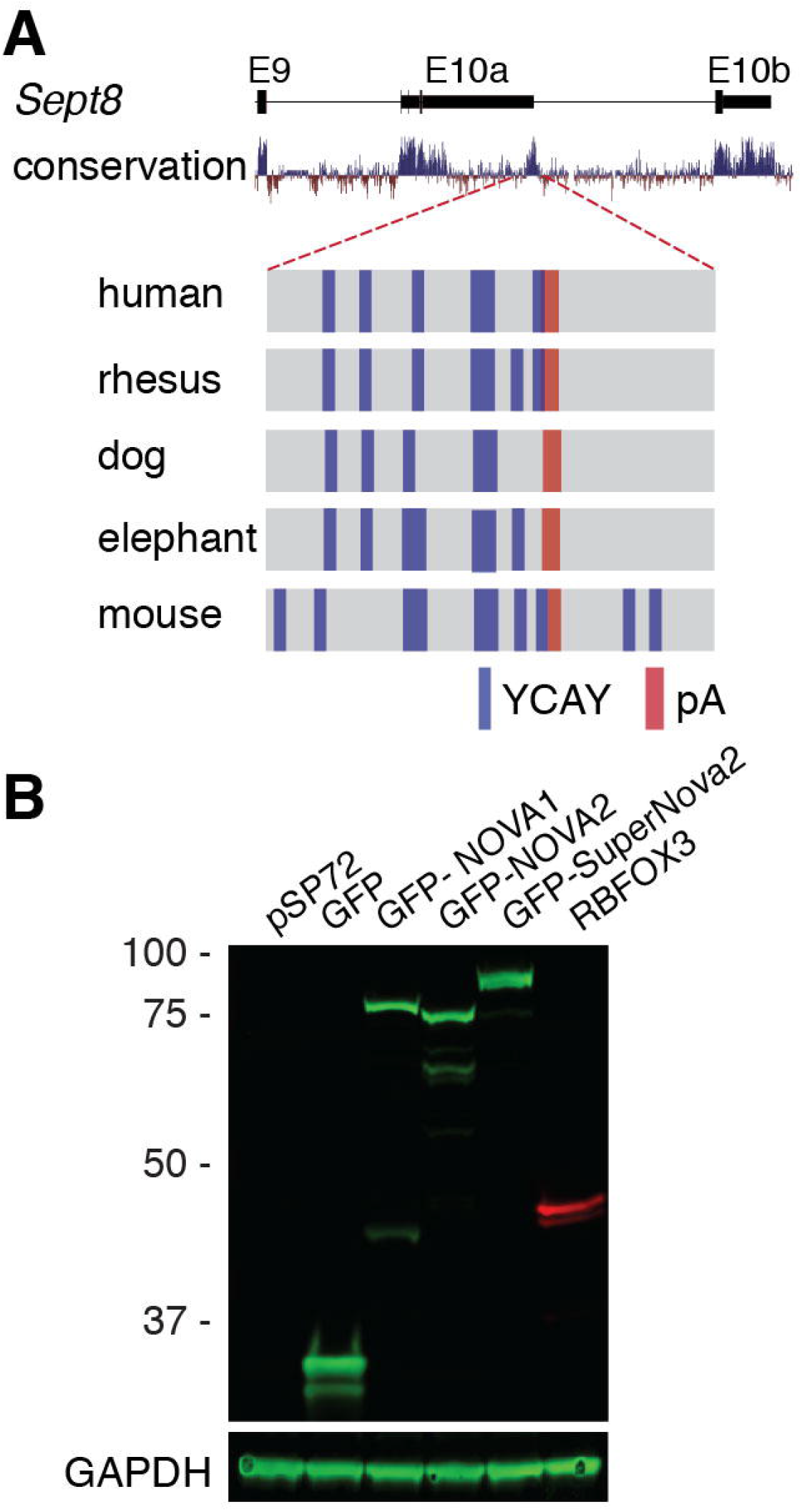

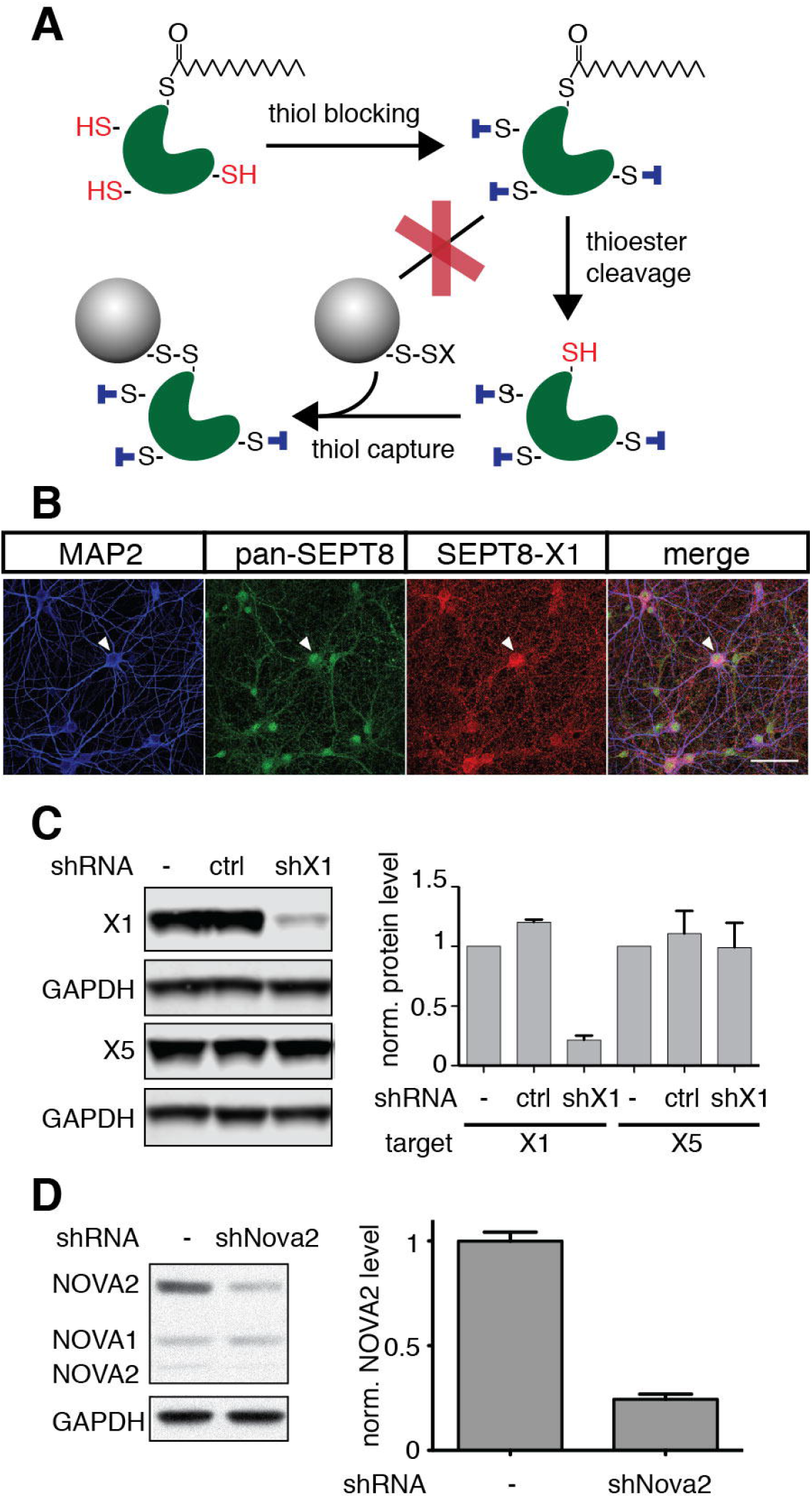

## References

1. Nelson SB, Sugino K, Hempel CM. The problem of neuronal cell types: a physiological genomics approach. Trends Neurosci. 2006;29:339–45.

2. Hobert O, Carrera I, Stefanakis N. The molecular and gene regulatory signature of a neuron. Trends Neurosci. 2010;33:435–45.

3. Fishell G, Heintz N. The neuron identity problem: form meets function. Neuron. 2013;80:602–12.

4. Darnell RB. RNA protein interaction in neurons. Annu. Rev. Neurosci. 2013;36:243–70.

5. Buckanovich RJ, Posner JB, Darnell RB. Nova, the paraneoplastic Ri antigen, is homologous to an RNA-binding protein and is specifically expressed in the developing motor system. Neuron. 1993;11:657–72.

6. Taliaferro JM, Vidaki M, Oliveira R, Olson S, Zhan L, Saxena T, et al. Distal Alternative Last Exons Localize mRNAs to Neural Projections. Molecular Cell. 2016;61:821–33.

7. Zhang X, Chen MH, Wu X, Kodani A, Fan J, Doan R, et al. Cell-Type-Specific Alternative Splicing Governs Cell Fate in the Developing Cerebral Cortex. CELL. 2016;166:1147–1162.e15.

8. Zhang Y, Chen K, Sloan SA, Bennett ML, Scholze AR, O’Keeffe S, et al. An RNA-sequencing transcriptome and splicing database of glia, neurons, and vascular cells of the cerebral cortex. J. Neurosci. 2014;34:11929–47.

9. Cahoy JD, Emery B, Kaushal A, Foo LC, Zamanian JL, Christopherson KS, et al. A transcriptome database for astrocytes, neurons, and oligodendrocytes: a new resource for understanding brain development and function. J. Neurosci. 2008;28:264–78.

10. Sanz E, Yang L, Su T, Morris DR, McKnight GS, Amieux PS. Cell-type-specific isolation of ribosome-associated mRNA from complex tissues. Proc. Natl. Acad. Sci. U.S.A. 2009;106:13939–44.

11. Heiman M, Kulicke R, Fenster RJ, Greengard P, Heintz N. Cell type-specific mRNA purification by translating ribosome affinity purification (TRAP). Nat Protoc. 2014;9:1282–91.

12. Heiman M, Schaefer A, Gong S, Peterson JD, Day M, Ramsey KE, et al. A translational profiling approach for the molecular characterization of CNS cell types. CELL. 2008;135:738–48.

13. Doyle JP, Dougherty JD, Heiman M, Schmidt EF, Stevens TR, Ma G, et al. Application of a translational profiling approach for the comparative analysis of CNS cell types. CELL. 2008;135:749–62.

14. Jensen KB, Musunuru K, Lewis HA, Burley SK, Darnell RB. The tetranucleotide UCAY directs the specific recognition of RNA by the Nova K-homology 3 domain. Proc. Natl. Acad. Sci. U.S.A. 2000;97:5740–5.

15. Ule J, Ule A, Spencer J, Williams A, Hu J-S, Cline M, et al. Nova regulates brain-specific splicing to shape the synapse. Nat Genet. 2005;37:844–52.

16. Licatalosi DD, Mele A, Fak JJ, Ule J, Kayikci M, Chi SW, et al. HITS-CLIP yields genome-wide insights into brain alternative RNA processing. Nature. 2008;456:464–9.

17. Ule J, Stefani G, Mele A, Ruggiu M, Wang X, Taneri B, et al. An RNA map predicting Nova-dependent splicing regulation. Nature. 2006;444:580–6.

18. Moore MJ, Zhang C, Gantman EC, Mele A, Darnell JC, Darnell RB. Mapping Argonaute and conventional RNA-binding protein interactions with RNA at single-nucleotide resolution using HITS-CLIP and CIMS analysis. Nat Protoc. 2014;9:263–93.

19. Darnell RB. HITS-CLIP: panoramic views of protein-RNA regulation in living cells. WIREs RNA. 2010;1:266–86.

20. Zhang C, Frias MA, Mele A, Ruggiu M, Eom T, Marney CB, et al. Integrative modeling defines the Nova splicing-regulatory network and its combinatorial controls. Science. 2010;329:439–43.

21. Ruggiu M, Herbst R, Kim N, Jevsek M, Fak JJ, Mann MA, et al. Rescuing Z+ agrin splicing in Nova null mice restores synapse formation and unmasks a physiologic defect in motor neuron firing. Proc. Natl. Acad. Sci. U.S.A. 2009;106:3513–8.

22. Huang CS, Shi S-H, Ule J, Ruggiu M, Barker LA, Darnell RB, et al. Common Molecular Pathways Mediate Long-Term Potentiation of Synaptic Excitation and Slow Synaptic Inhibition. CELL. 2005;123:105–18.

23. Yano M, Hayakawa-Yano Y, Mele A, Darnell RB. Nova2 regulates neuronal migration through an RNA switch in disabled-1 signaling. Neuron. 2010;66:848–58.

24. Saito Y, Miranda-Rottmann S, Ruggiu M, Park CY, Fak JJ, Zhong R, et al. NOVA2-mediated RNA regulation is required for axonal pathfinding during development. Elife. 2016;5.

25. Yang YY, Yin GL, Darnell RB. The neuronal RNA-binding protein Nova-2 is implicated as the autoantigen targeted in POMA patients with dementia. Proc. Natl. Acad. Sci. U.S.A. 1998;95:13254–9.

26. Charizanis K, Lee KY, Batra R, Goodwin M, Zhang C, Yuan Y, et al., Muscleblind-like 2-Mediated Alternative Splicing in the Developing Brain and Dysregulation in Myotonic Dystrophy. Neuron. 2012;75:437–50.

27. Shah A, Qian Y, Weyn-Vanhentenryck SM, Zhang C. CLIP Tool Kit (CTK): a flexible and robust pipeline to analyze CLIP sequencing data. Bioinformatics. 2017;33:566–7.

28. Gong S, Doughty M, Harbaugh CR, Cummins A, Hatten ME, Heintz N, et al. Targeting Cre recombinase to specific neuron populations with bacterial artificial chromosome constructs. J. Neurosci. 2007;27:9817–23.

29. Robinson MD, McCarthy DJ, Smyth GK. edgeR: a Bioconductor package for differential expression analysis of digital gene expression data. Bioinformatics. 2010;26:139–40.

30. Enjin A, Rabe N, Nakanishi ST, Vallstedt A, Gezelius H, Memic F, et al. Identification of novel spinal cholinergic genetic subtypes disclose Chodl and Pitx2 as markers for fast motor neurons and partition cells. J. Comp. Neurol. 2010;518:2284–304.

31. Kodama T, Guerrero S, Shin M, Moghadam S, Faulstich M, Lac du S. Neuronal classification and marker gene identification via single-cell expression profiling of brainstem vestibular neurons subserving cerebellar learning. J. Neurosci. 2012;32:7819–31.

32. Vullhorst D, Neddens J, Karavanova I, Tricoire L, Petralia RS, McBain CJ, et al. Selective expression of ErbB4 in interneurons, but not pyramidal cells, of the rodent hippocampus. J. Neurosci. 2009;29:12255–64.

33. Neddens J, Fish KN, Tricoire L, Vullhorst D, Shamir A, Chung W, et al. Conserved interneuron-specific ErbB4 expression in frontal cortex of rodents, monkeys, and humans: implications for schizophrenia. Biol. Psychiatry. 2011;70:636–45.

34. Rexed B. The cytoarchitectonic organization of the spinal cord in the cat. J. Comp. Neurol. 1952;96:414–95.

35. Henry AM, Hohmann JG. High-resolution gene expression atlases for adult and developing mouse brain and spinal cord. Mamm. Genome. 2012;23:539–49.

36. Bandyopadhyay U, Cotney J, Nagy M, Oh S, Leng J, Mahajan M, et al. RNA-Seq profiling of spinal cord motor neurons from a presymptomatic SOD1 ALS mouse. PLoS ONE. 2013;8:e53575.

37. Saarikangas J, Kourdougli N, Senju Y, Chazal G, Segerstråle M, Minkeviciene R, et al. MIM-Induced Membrane Bending Promotes Dendritic Spine Initiation. Dev. Cell. 2015;33:644–59.

38. Yu N, Signorile L, Basu S, Ottema S, Lebbink JHG, Leslie K, et al. Isolation of Functional Tubulin Dimers and of Tubulin-Associated Proteins from Mammalian Cells. Curr. Biol. 2016;26:1728–36.

39. Lee K-H, Lee JS, Lee D, Seog D-H, Lytton J, Ho W-K, et al. KIF21A-mediated axonal transport and selective endocytosis underlie the polarized targeting of NCKX2. J. Neurosci. 2012;32:4102–17.

40. Baines AJ, Lu H-C, Bennett PM. The Protein 4.1 family: hub proteins in animals for organizing membrane proteins. Biochim. Biophys. Acta. 2014;1838:605–19.

41. Al-Bassam J, Chang F. Regulation of microtubule dynamics by TOG-domain proteins XMAP215/Dis1 and CLASP. Trends Cell Biol. 2011;21:604–14.

42. Galjart N. CLIPs and CLASPs and cellular dynamics. Nat. Rev. Mol. Cell Biol. 2005;6:487–98.

43. Zhang Y, Zhang X-F, Fleming MR, Amiri A, El-Hassar L, Surguchev AA, et al. Kv3.3 Channels Bind Hax-1 and Arp2/3 to Assemble a Stable Local Actin Network that Regulates Channel Gating. CELL. 2016;165:434–48.

44. Kirsch J, Betz H. The postsynaptic localization of the glycine receptor-associated protein gephyrin is regulated by the cytoskeleton. J. Neurosci. 1995;15:4148–56.

45. Mattila PK, Salminen M, Yamashiro T, Lappalainen P. Mouse MIM, a tissue-specific regulator of cytoskeletal dynamics, interacts with ATP-actin monomers through its C-terminal WH2 domain. J. Biol. Chem. 2003;278:8452–9.

46. Glassmann A, Molly S, Surchev L, Nazwar TA, Holst M, Hartmann W, et al. Developmental expression and differentiation-related neuron-specific splicing of metastasis suppressor 1 (Mtss1) in normal and transformed cerebellar cells. BMC Dev. Biol. 2007;7:111.

47. Sistig T, Lang F, Wrobel S, Baader SL, Schilling K, Eiberger B. Mtss1 promotes maturation and maintenance of cerebellar neurons via splice variant-specific effects. Brain Struct Funct. 2017.

48. Goldman-Wohl DS, Chan E, Baird D, Heintz N. Kv3.3b: a novel Shaw type potassium channel expressed in terminally differentiated cerebellar Purkinje cells and deep cerebellar nuclei. J. Neurosci. 1994;14:511–22.

49. Brooke RE, Atkinson L, Edwards I, Parson SH, Deuchars J. Immunohistochemical localisation of the voltage gated potassium ion channel subunit Kv3.3 in the rat medulla oblongata and thoracic spinal cord. Brain Res. 2006;1070:101–15.

50. Gladfelter AS, Bose I, Zyla TR, Bardes ESG, Lew DJ. Septin ring assembly involves cycles of GTP loading and hydrolysis by Cdc42p. J. Cell Biol. 2002;156:315–26.

51. Sadian Y, Gatsogiannis C, Patasi C, Hofnagel O, Goody RS, Farkasovský M, et al. The role of Cdc42 and Gic1 in the regulation of septin filament formation and dissociation. Elife. 2013;2:e01085.

52. Kinoshita M. Assembly of mammalian septins. J. Biochem. 2003;134:491–6.

53. Weirich CS, Erzberger JP, Barral Y. The septin family of GTPases: architecture and dynamics. Nat. Rev. Mol. Cell Biol. 2008;9:478–89.

54. Mostowy S, Cossart P. Septins: the fourth component of the cytoskeleton. Nat. Rev. Mol. Cell Biol. 2012;13:183–94.

55. Tada T, Simonetta A, Batterton M, Kinoshita M, Edbauer D, Sheng M. Role of Septin cytoskeleton in spine morphogenesis and dendrite development in neurons. Curr. Biol. 2007;17:1752–8.

56. Xie Y, Vessey JP, Konecna A, Dahm R, Macchi P, Kiebler MA. The GTP-binding protein Septin 7 is critical for dendrite branching and dendritic-spine morphology. Curr. Biol. 2007;17:1746–51.

57. Hu J, Bai X, Bowen JR, Dolat L, Korobova F, Yu W, et al. Septin-driven coordination of actin and microtubule remodeling regulates the collateral branching of axons. Curr. Biol. 2012;22:1109–15.

58. Kang R, Wan J, Arstikaitis P, Takahashi H, Huang K, Bailey AO, et al. Neural palmitoyl-proteomics reveals dynamic synaptic palmitoylation. Nature. 2008;456:904–9.

59. Wan J, Savas JN, Roth AF, Sanders SS, Singaraja RR, Hayden MR, et al. Tracking brain palmitoylation change: predominance of glial change in a mouse model of Huntington’s disease. Chem. Biol. 2013;20:1421–34.

60. Xie Y, Zheng Y, Li H, Luo X, He Z, Cao S, et al. GPS-Lipid: a robust tool for the prediction of multiple lipid modification sites. Sci Rep. 2016;6:28249.

61. Forrester MT, Hess DT, Thompson JW, Hultman R, Moseley MA, Stamler JS, et al. Site-specific analysis of protein S-acylation by resin-assisted capture. J. Lipid Res. 2011;52:393–8.

62. Antinone SE, Ghadge GD, Ostrow LW, Roos RP, Green WN. S-acylation of SOD1, CCS, and a stable SOD1-CCS heterodimer in human spinal cords from ALS and nonALS subjects. Sci Rep. 2017;7:41141.

63. Gauthier-Campbell C, Bredt DS, Murphy TH, El-Husseini AE-D. Regulation of dendritic branching and filopodia formation in hippocampal neurons by specific acylated protein motifs. Mol. Biol. Cell. 2004;15:2205–17.

64. Abdel-Maguid TE, Bowsher D. Alpha- and gamma-motoneurons in the adult human spinal cord and somatic cranial nerve nuclei: the significance of dendroachitectonics studied by the Golgi method. J. Comp. Neurol. 1979;186:259–69.

65. Paxinos G. The Human Nervous System. Academic Press; 2012.

66. Martin E, Cazenave W, Cattaert D, Branchereau P. Embryonic alteration of motoneuronal morphology induces hyperexcitability in the mouse model of amyotrophic lateral sclerosis. Neurobiol. Dis. 2013;54:116–26.

67. Kato T, Hirano A, Donnenfeld H. A Golgi study of the large anterior horn cells of the lumbar cords in normal spinal cords and in amyotrophic lateral sclerosis. Acta Neuropathol. 1987;75:34–40.

68. Fu X-D, Ares M. Context-dependent control of alternative splicing by RNA-binding proteins. Nature Publishing Group. 2014;15:689–701.

69. Polydorides AD, Okano HJ, Yang YY, Stefani G, Darnell RB. A brain-enriched polypyrimidine tract-binding protein antagonizes the ability of Nova to regulate neuron-specific alternative splicing. Proc. Natl. Acad. Sci. U.S.A. 2000;97:6350–5.

70. Buckanovich RJ, Yang YY, Darnell RB. The onconeural antigen Nova-1 is a neuron-specific RNA-binding protein, the activity of which is inhibited by paraneoplastic antibodies. J. Neurosci. 1996;16:1114–22.

71. Schaefer A, Im H-I, Venø MT, Fowler CD, Min A, Intrator A, et al. Argonaute 2 in dopamine 2 receptor-expressing neurons regulates cocaine addiction. J. Exp. Med. 2010;207:1843–51.

72. Tan CL, Plotkin JL, Venø MT, Schimmelmann von M, Feinberg P, Mann S, et al. MicroRNA-128 governs neuronal excitability and motor behavior in mice. Science. 2013;342:1254–8.

73. Bahn JH, Ahn J, Lin X, Zhang Q, Lee J-H, Civelek M, et al. Genomic analysis of ADAR1 binding and its involvement in multiple RNA processing pathways. Nat Commun. 2015;6:6355.

74. Beltran M, Yates CM, Skalska L, Dawson M, Reis FP, Viiri K, et al. The interaction of PRC2 with RNA or chromatin is mutually antagonistic. Genome Research. 2016;26:896–907.

75. Hwang H-W, Park CY, Goodarzi H, Fak JJ, Mele A, Moore MJ, et al. PAPERCLIP Identifies MicroRNA Targets and a Role of CstF64/64tau in Promoting Non-canonical poly(A) Site Usage. Cell Reports. 2016;15:423–35.

76. Pramatarova A, Laganière J, Roussel J, Brisebois K, Rouleau GA. Neuron-specific expression of mutant superoxide dismutase 1 in transgenic mice does not lead to motor impairment. J. Neurosci. 2001;21:3369–74.

77. Lino MM, Schneider C, Caroni P. Accumulation of SOD1 mutants in postnatal motoneurons does not cause motoneuron pathology or motoneuron disease. J. Neurosci. 2002;22:4825–32.

78. Boillée S, Vande Velde C, Cleveland DW. ALS: a disease of motor neurons and their nonneuronal neighbors. Neuron. 2006;52:39–59.

79. Phatnani HP, Guarnieri P, Friedman BA, Carrasco MA, Muratet M, O’Keeffe S, et al. Intricate interplay between astrocytes and motor neurons in ALS. Proc. Natl. Acad. Sci. U.S.A. 2013;110:E756–65.

80. Weyn-Vanhentenryck SM, Mele A, Yan Q, Sun S, Farny N, Zhang Z, et al. HITS-CLIP and integrative modeling define the Rbfox splicing-regulatory network linked to brain development and autism. Cell Reports. 2014;6:1139–52.

81. Gong S, Yang XW, Li C, Heintz N. Highly efficient modification of bacterial artificial chromosomes (BACs) using novel shuttle vectors containing the R6Kgamma origin of replication. Genome Research. 2002;12:1992–8.

82. Zhang C, Darnell RB. Mapping in vivo protein-RNA interactions at single-nucleotide resolution from HITS-CLIP data. Nat. Biotechnol. 2011;29:607–14.

83. Darnell JC, Van Driesche SJ, Zhang C, Hung KYS, Mele A, Fraser CE, et al. FMRP stalls ribosomal translocation on mRNAs linked to synaptic function and autism. CELL. 2011;146:247–61.

84. Wu J, Anczuków O, Krainer AR, Zhang MQ, Zhang C. OLego: fast and sensitive mapping of spliced mRNA-Seq reads using small seeds. Nucleic Acids Research. 2013;41:5149–63.

85. Eden E, Navon R, Steinfeld I, Lipson D, Yakhini Z. GOrilla: a tool for discovery and visualization of enriched GO terms in ranked gene lists. BMC Bioinformatics. 2009;10:48.

